# The eIF4A2 negative regulator of mRNA translation promotes extracellular matrix deposition to accelerate hepatocellular carcinoma initiation

**DOI:** 10.1101/2023.08.16.553544

**Authors:** Madeleine Moore, Luis Pardo-Fernandez, Louise Mitchell, Tobias Schmidt, Joseph A Waldron, Stephanie May, Miryam Muller, Rachael C. L. Smith, Douglas Strathdee, Sheila Bryson, Kelly Hodge, Sergio Lilla, Ania Wilczynska, Lynn McGarry, Sarah Gillen, Ruban Peter-Durairaj, Georgios Kanellos, Colin Nixon, Sara Zanivan, Owen J. Sansom, Thomas G. Bird, Martin Bushell, Jim C. Norman

**Affiliations:** Cancer Research UK Scotland Institute, Glasgow, G61 1BD, UK; MRC Centre for Inflammation Research, The Queen’s Medical Research Institute, University of Edinburgh, EH16 4TJ, UK; School of Cancer Sciences, University of Glasgow, Glasgow, G61 1QH, UK

## Abstract

Increased protein synthesis supports growth of established tumours. However, how mRNA translation contributes to early tumorigenesis remains unclear. Here we show that following oncogene activation, hepatocytes enter a non-proliferative/senescent-like phase characterized by α5β1 integrin-dependent deposition of fibronectin-rich extracellular matrix (ECM) niches. These niches then promote exit from oncogene-induced senescence to permit progression to proliferating hepatocellular carcinoma (HCC). Removal of eIF4A2, a negative regulator of mRNA translation, boosts the synthesis of membrane/secretory proteins which drives a compensatory increase in the turnover/degradation of membrane proteins including α5β1 integrin. This increased membrane protein degradation, in turn, compromises generation of ECM-rich tumour initiation niches, senescence-exit and progression to proliferating HCC. Consistently, pharmacological inhibition of mRNA translation following eIF4A2 loss restores ECM deposition and reinstates HCC progression. Thus, although inhibition of protein synthesis may be an effective way to reduce tumour biomass and the growth of established tumours, our results highlight how agents which reduce mRNA translation, if administered during early tumorigenesis, may awaken senescent cells and promote tumour progression.

## Main

Several of cancer’s aggressive characteristics depend on the ability of oncogenic signaling to re-program mRNA-translation^1, 2^. The initiation of mRNA-translation is a critical event in selection of mRNAs for translation and the eIF4F complex is the master regulator of this process ^3, 4^). Certain signaling pathways which are dysregulated following oncogene activation – particularly those culminating in elevated TORC1 signaling – converge on eIF4F in a way that favours synthesis of proteins promoting tumour cell proliferation, invasion, and metastasis^1, 5, 6^. These observations have inspired initiatives to inhibit signaling converging on the eIF4F complex as an attractive strategy towards reversing both the uncontrolled growth and the invasive/metastatic behaviour of cancers. This has proved productive as agents designed to inhibit the translational initiation machinery downstream of oncogenic signaling lead, not only to reduced tumour growth, but to alterations to the tumour microenvironment, which may increase the efficacy of anti-tumour immunotherapy^7–12^. Thus, a growing literature is beginning to highlight the importance of translation initiation and, specifically eIF4F, in influencing both tumour cell autonomous patterns of behaviour, such as growth and invasiveness, and the more complex landscape of the local and systemic tumour microenvironment. The eIF4A mRNA helicase is a key component of the eIF4F complex, and its principal role is to load 40S ribosomes onto mRNA and then contribute to the unwinding of structure within the mRNA, allowing ribosomes to scan the 5’-UTR, find a start codon and initiate translation^6^. There are two eIF4A paralogue proteins (eIF4A1 and eIF4A2) and, although both possess RNA helicase activity, eIF4A1 associates with the eIF4F complex to activate translation initiation whilst eIF4A2 exhibits more complex behaviour. eIF4A2 can recruit the CCR4-NOT complex to repress translation and promote mRNA degradation^13, 14^, but in the absence of eIF4A1, it has been shown to bind the eIF4F complex, possibly to promote translation. Importantly, eIF4A paralogues exhibit distinct tissue expression patterns, with high eIF4A1 levels being associated with proliferative cells. For example, eIF4A1 expression is high in intestinal crypts where epithelial proliferation occurs, whilst eIF4A2 is expressed throughout the colonic epithelial axis and is particularly abundant in the secretory cells near the top of the crypt [see accompanying manuscript]. Consistently, eIF4A1 but not eIF4A2 deletion in the gut opposes hyperproliferative phenotypes, suggesting that eIF4A1 and eIF4A2 may support proliferative and secretory translational landscapes respectively. Moreover, the CCR4-NOT complex – which can be associated with eIF4A2 - has been shown to influence translational selectivity in a way that may favour secretory phenotypes. Indeed, translation of mRNAs encoding secretory proteins, such as extracellular matrix (ECM) components, are differentially affected following CCR4-NOT disruption^15^.

Carcinoma initiation involves complex interplay between acquisition of oncogenic mutations in epithelial cells, and establishment of a supportive microenvironment. Liver cancer or hepatocellular carcinoma (HCC) is a blueprint for this paradigm; its initiation being characterized by accumulation of oncogenic mutations typically alongside conditions such as non-alcoholic steatohepatitis (NASH)-with-fibrosis and liver cirrhosis, which are characterized by chronic deposition of fibrotic ECM proteins^16, 17^. Despite advances in our understanding of how eIF4F maintains cancer growth and aggression, little is known as to how protein synthesis landscapes influence tumour initiation and subsequently sculpt the microenvironment in early tumorigenesis. To address this, we have specifically targeted the translation initiation machinery in genetically engineered mouse models (GEMMs) which recapitulate early tumorigenesis and progression of HCC. We identified that modulation of translation initiation by eIF4A2 – a factor which we find to selectively suppress translation of membrane and secreted components – is required for the establishment of ECM-rich niches which promote progression of hepatocytes from a state of oncogene-induced senescence to rapidly proliferating and end-stage HCC.

## Results

### eIF4A2 enables α5β1 integrin-dependent ECM deposition in early tumorigenesis to license progression to aggressive HCC

To model initiation and progression of HCC *in vivo*, we used mice carrying floxed alleles of constitutively active β-catenin (*Ctnnb1^ex3flox/WT^)* and additional copies of the human *MYC* oncogene (*R26*^LSL-MYC/LSL-MYC^). Expression of these oncogenes was initiated by injecting mice with AAV8-TBG-Cre at a titre that achieves recombination in fewer than 5% of hepatocytes; the tumour cell-of-origin for HCC^18, 19^ (Fig. 1A). Staining for glutamine synthetase (GS) – a WNT target gene - indicated functional activation of WNT signaling downstream of active β-catenin in the recombined cell population (Fig. 1B). Within 5 days of oncogene induction many of the recombined, oncogene-expressing cells were positive for Ki67, indicating that they had started to proliferate (Fig. 1A-B). Following this proliferative burst (30 days after AAV-TBG-Cre injection), Ki67-positive cells were scarce. Rather, we observed the presence of a population of enlarged hepatocytes which were Ki67 negative and p21-positive which we term pre-malignant oncogene-induced senescent hepatocytes (Fig. 1A-C; S1A). Following this period of senescence, a subset of the GS-positive hepatocytes resumed proliferation. Numerous proliferative lesions/tumour nodules were visible 90 days following AAV-TBG-Cre injection (Fig. 1A-B&D) which continued to develop into end-stage HCC leading to death at approximately 135 days (Fig. 1B&E).

**Fig. 1.**
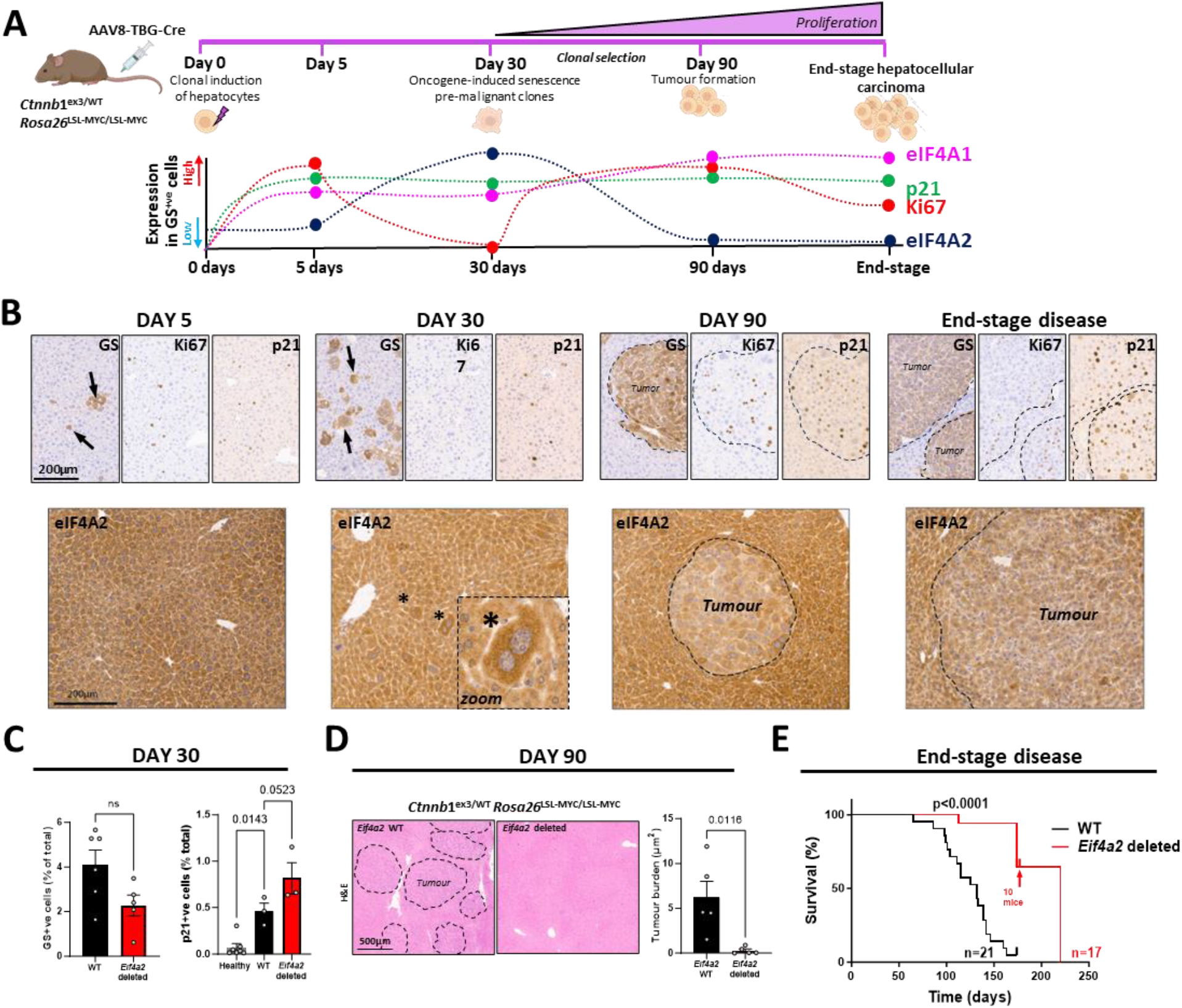
eIF4A2 enables early senescent lesions to progress to aggressive HCC. *Ctnnb*1^ex3/WT^; *Rosa26*^LSL-MYC/LSL-MYC^ mice that were either *Eif4a2*^WT/WT^ *or Eif4a2*^fl/fl^ were injected (i.v.) with AAV-TBG-Cre (6.4 X 10^8^ genome copies (GC) per mouse; a titre sufficient to evoke recombination in ≈5% hepatocytes). Mice were sacrificed 5 (B), 30 (**B, C**), or 90 (**B, D**) days following this or at tumour endpoint (**B, E**; n=21 *Eif4a1/2*^WT/WT^ mice and n=7 *Eif4a2*^fl/fl^ mice). An additional 10 *Eif4a2*^fl/fl^ mice were sacrificed at 175 days following AAV-TBG-Cre injection, as annotated on the figure (**E**). Glutamine synthetase (GS; an indicator of activated β-catenin signaling), p21, Ki67, eIF4A1 and eIF4A2 were visualized in *Ctnnb*1^ex3/WT^; *Rosa26*^LSL-MYC/LSL-MYC^ mice in zone 2 of the liver by immunohistochemistry (**B**) and levels of these are summarized in the schematic in (**A**). GS-positive induced hepatocytes are indicated by arrows in (B). Induced hepatocytes with increased levels of eIF4A2 are denoted by asterisks in and highlighted in the ‘zoom’ in **(B).** GS- and p21-positive cells were quantified 30 days following AAV-TBG-Cre injection using high content analysis and immunohistochemistry respectively (**C**). Tumour burden was determined by H&E staining (**D**). All conditions depicted in figure 1 correspond to *Ctnnb*1^ex3/WT^; *Rosa26*^LSL-MYC/LSL-MYC^ mice except for ‘Healthy’ mice where *Ctnnb*1^ex3/WT^; *Rosa26*^LSL-MYC/LSL-MYC^ mice were injected with AAV TGB Null virus (6.4 X 10^8^ genome copies (GC) per mouse). Each point in the graphs in **(C, D**) represents quantification of staining from an individual mouse liver – values are mean±SEM, statistical significance is indicated by *p* value, data were analyzed by one-way ANOVA. The statistical test used for the Kaplan-Maier analyses in (**E**) was Log rank (Mantel-Cox), and the cohort sizes used to generate these are displayed on the graph.

We monitored expression of the two eIF4A paralogues throughout HCC initiation and progression and found that 30 days following AAV-TBG-Cre injection, both eIF4A1 and eIF4A2 were upregulated in GS-positive hepatocytes (Fig. 1A,B; S1B,C). The subsequent transition to proliferating lesions was accompanied by reduced levels of eIF4A2, whilst eIF4A1 expression was maintained until tumour endpoint (Fig. 1A-B; S1C). We hypothesized that the transient upregulation of eIF4A2 observed 30 days following oncogene activation may indicate a role for this translation repressor in early tumorigenesis. To investigate this, we generated mice with floxed alleles of *Eif4a1* and *Eif4a2* (see accompanying manuscript) and crossed them into the β-catenin/Myc-driven GEMM of HCC. Knockout of *Eif4a2* (Fig. S1D), strongly inhibited progression of these lesions to proliferative HCC (Fig. 1D) and significantly extended survival of the mice (Fig. 1E). By contrast, *Eif4a1* knockout was ineffective in this regard (Fig. S1E). *Eif4a2* deletion, however, did not compromise the GS/p21-positive population of pre-malignant senescent hepatocytes observed 30 days following AAV-TBG-Cre injection (Fig. 1C), suggesting that this paralogue may contribute to the progression from oncogene-induced senescence to proliferative HCC.

HCC initiation involves interplay between oncogene activation and alteration to the liver microenvironment^17^. Accumulation of fibrillar ECM components, such as fibronectin (FN), is strongly associated with the genesis of HCC, and α5 integrin (ITGA5).

The α-chain of the α5β1 integrin heterodimer responsible for efficient FN polymerization/deposition^20^, is significantly upregulated in GS-positive hepatocytes 30 days following injection of AAV-TBG-Cre (Fig. 2A).

**Fig. 2.**
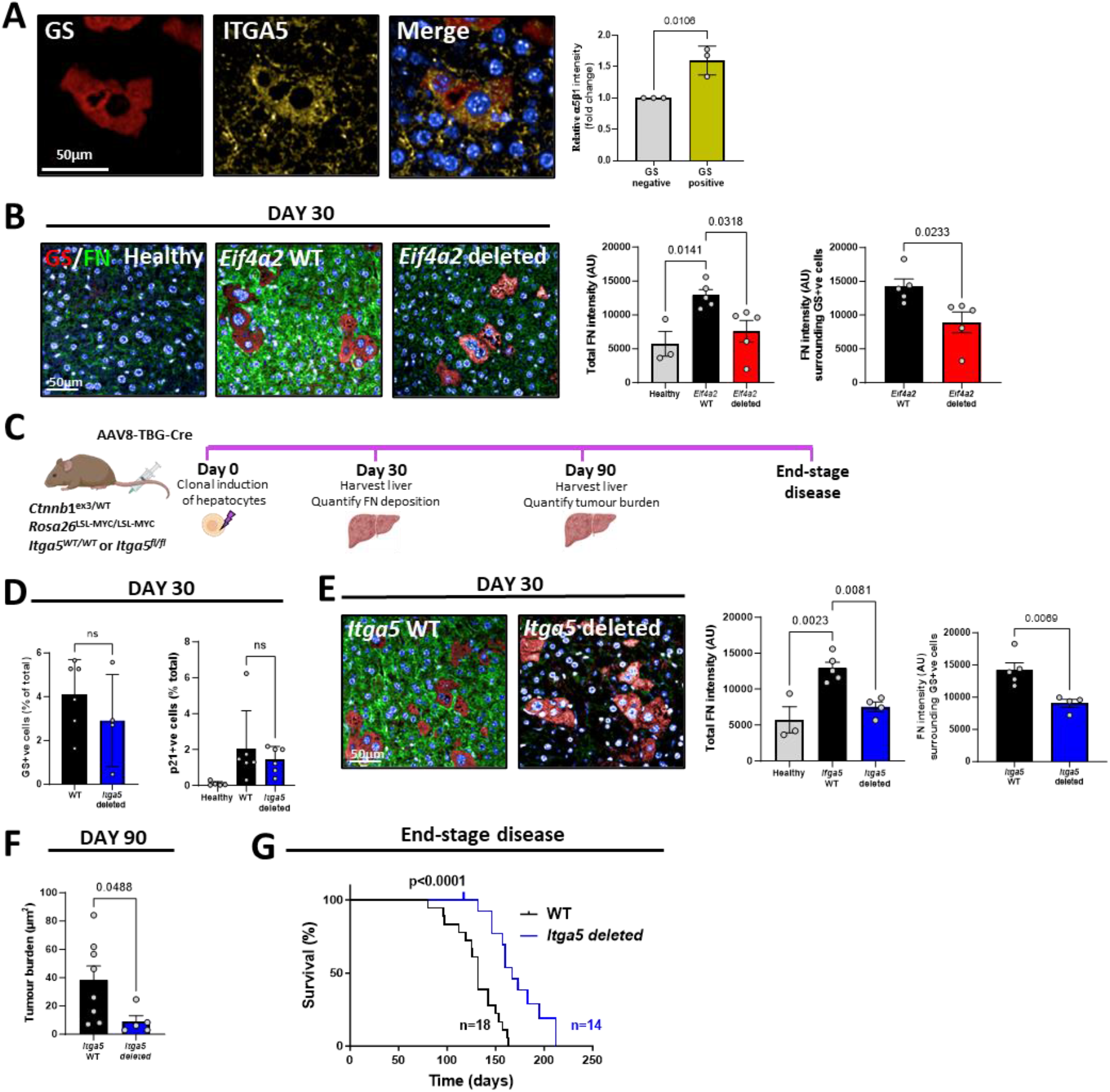
eIF4A2 controls α5β1 integrin-dependent fibronectin deposition to drive HCC progression. *Ctnnb*1^ex3/WT^; *Rosa26*^LSL-MYC/LSL-MYC^ mice that were either *Eif4a2*^WT/W^, *Eif4a2*^fl/fl^, *Itga5*^WT/WT^ or *Itga5*^fl/fl^ were injected (i.v.) with AAV-TBG-Cre (6.4 X 10^8^ genome copies (GC) per mouse; a titre sufficient to evoke recombination in ≈5% hepatocytes). Mice were sacrificed 30 (**A,B,D,E**), or 90 (**F**) days following this or at tumor endpoint (**G**). GS (red), ITGA5 (yellow), FN (green) and p21 were visualized and quantified in zone 2 of the liver. Quantification of FN and IGTA5 levels were by immunofluorescence followed by high content analysis (**A,B,E**). Tumour burden was determined by H&E staining (**F**). Each point in the graphs in (**A,B,D,E,F**) represents quantification of staining from an individual mouse liver – values are mean±SEM, statistical significance is indicated by *p* value, data were analyzed by one-way ANOVA. The statistical test used for the Kaplan-Maier analysis in (**G**) was Log rank (Mantel-Cox), and the cohort sizes used to generate these are displayed on the graph.

Consistently, activation of β-Catenin/Myc led to increased FN deposition in the vicinity of GS-positive hepatocytes and more widely in the liver, and this was opposed by *Eif4a2* knockout (Fig. 2B). Hepatocyte-specific deletion of *Itga5*^20^, whilst not affecting the ability of β-catenin/MYC activation to drive the appearance of p21/GS-positive hepatocytes (Fig. 2C,D), prevented FN deposition associated with these cells (Fig. 2E) and retarded their progression to proliferative HCC, thus significantly extending survival (Fig. 2F,G).

### eIF4A2 moderates translation to reduce membrane/secretory protein turnover and enable ECM deposition

Hepatocytes respond to genetic and toxic insults by adopting a state characterized by a senescence-associated secretory phenotype (SASP) ^21–24^. As well as driving stellate cell activation (via soluble factors, such as TGFβ), which can lead to extensive deposition of fibrotic ECM, the hepatocyte SASP itself contains ECM components, including FN^25^. Our finding that hepatocytes must express α5β1 integrin to enable FN polymerization following oncogene induction (Fig. 2E), suggests that these epithelial cells make direct contributions to fibrillar ECM deposition. We injected mice bearing floxed alleles of murine double minute (*Mdm2*) with AAV8-TBG-Cre at a titre that achieves recombination and senescence activation in more than 90% of hepatocytes after 4 days^21^ (Fig. 3A). Here, we detected activation of the p53/p21 axis leading to widespread hepatocyte senescence and extensive FN deposition (Fig. 3B). Combined hepatocyte-specific *Eif4a2* (but not *Eif4a1*) (Fig. S2A) and *Mdm2* deletion reversed senescence-associated deposition of FN (Fig. 3B; Fig. S2B) but did not reduce implementation of the senescence phenotype (as evidenced by unchanged p21 expression) (Fig. 3B). Although hepatocyte-specific *Mdm2* deletion increased α- smooth muscle actin (αSMA)-positive/activated stellate cells in the liver, this was unaffected by *Eif4a2* knockout (Fig. 3B). This indicates that, whilst stellate cell activation occurring downstream of hepatocyte senescence is eIF4A2 independent, eIF4A2 expression by senescent hepatocytes is required for FN deposition by this epithelial cell type.

**Fig. 3.**
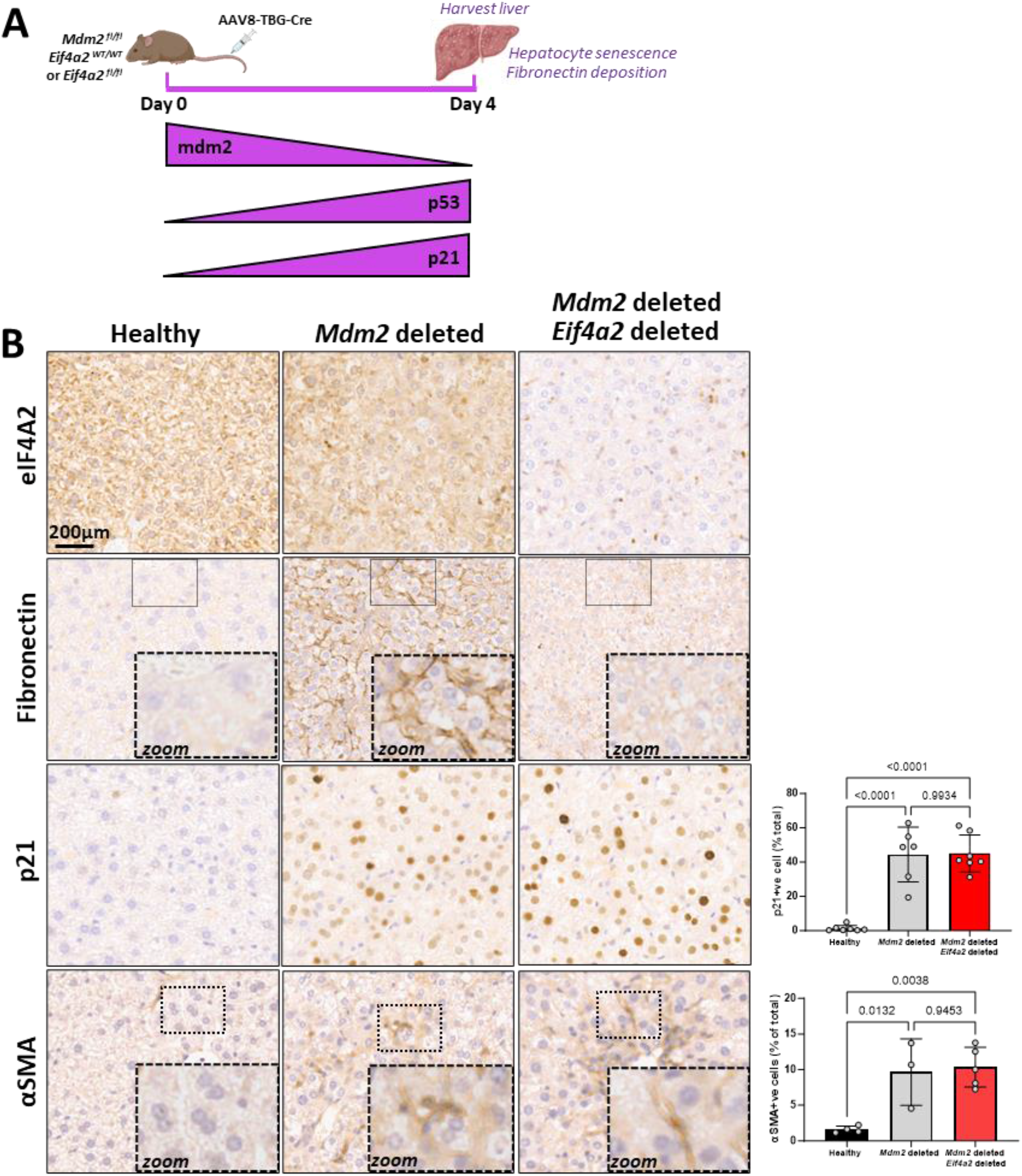
eIF4A2 is required for senescence-associated fibronectin deposition in the liver. *Mdm2*^fl/fl^ mice that were either *Eif4a2*^WT/WT^ or *Eif4a2*^fl/fl^ were injected with AAV-TBG-Cre (2 X 10^11^ GC per mouse; a titre sufficient to evoke recombination in >90% hepatocytes) and sacrificed 4 days later (**A**). *Mdm2*^fl/fl^ mice were injected with AAV-TGB-Null virus (2 X 10^11^ GC per mouse) to generate healthy control tissue. eIF4A2, fibronectin, p21, and α-smooth muscle actin (αSMA) were visualized in zone 2 of the liver using immunohistochemistry. Each point in the graphs in (**B**) represents data from an individual mouse liver. Values are mean±SEM, statistical significance is indicated by p values, data were analyzed by one-way ANOVA.

To understand how eIF4A2’s role as a translation regulator might enable FN deposition *in vivo*, we performed polysome profiling and found that *Eif4a2* deletion did not significantly influence the overall ratio of polysomes to sub-polysomes (Fig. S3A) in *Mdm2* knockout mouse livers. Nevertheless, RNAseq indicated that mRNAs classified as ‘external encapsulating structure’ – a category comprising secreted ECM components and plasma membrane receptors (including integrins) – were prominent amongst those displaying *increased* polysome association following *Eif4a2* deletion (Fig. S3B,C).

This finding is consistent with previous observations that knockdown of the eIF4A2-associated factor, CCR4-NOT, selectively influences translation of mRNAs encoding ECM components^15^. This *in vivo* observation presented a paradox in which *increased* synthesis of ECM components accompanied *decreased* ECM deposition. To resolve this, we needed to gain a comprehensive view of how *Eif4a2* deletion affects global protein degradation and synthesis landscapes.

To study eIF4A2’s role in controlling ECM synthesis and degradation, we used a model in which primary cultured fibroblasts enter senescence following inducible expression of oncogenic HRAS (Fig. 4A) ^26^. Oncogene-induced senescence was associated with increased levels of eIF4A2 (Fig. 4B), and elevated deposition of fibrillar FN which was significantly reduced following eIF4A2 loss (Fig. 4C), indicating that this *in vitro* model of senescence recapitulates the role played by eIF4A2 in ECM deposition *in vivo*. Pulsed-SILAC^27^, which quantifies incorporation of a pulse of labelled amino acids into newly synthesized protein on a global proteomic level, indicated that eIF4A1 and eIF4A2 had opposing effects on protein synthesis; with eIF4A1 and eIF4A2 acting to activate (Fig. S4) or repress (Fig. 4D) translation respectively. Notably, ECM-related proteins (including the ‘matrisome’) were prominent amongst the polypeptides whose synthesis was increased by eIF4A2 loss (Fig. 4D,F). This is consistent with our observations that *in vivo* deletion of *Eif4a2* in senescent hepatocytes increases polysome recruitment of mRNAs encoding ECM proteins (Fig. S3B,C).

**Fig. 4.**
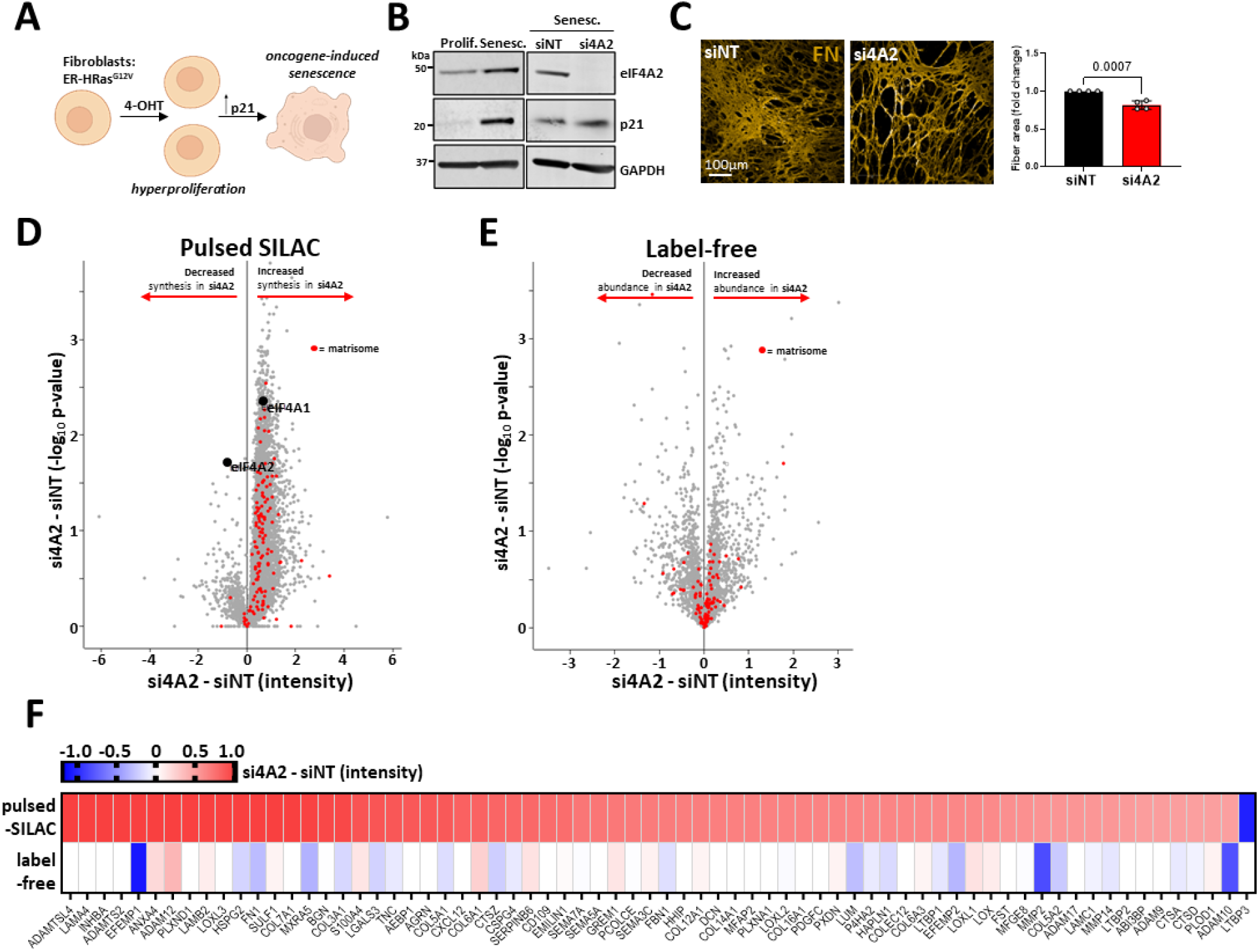
eIF4A2 represses translation of the matrisome. Primary cultured lung fibroblasts expressing the inducible ER-HRAS^G12V^ oncogene and transfected with either non-targeting (siNT) oligonucleotides or those targeting eIF4A2 (si4A2) were treated with 4-hydroxytamoxifen (4-OHT) to induce oncogene-induced senescence **(A)**. Expression of p21 and eIF4A2 was monitored by western blot **(B)**. Senescent cells were subsequently allowed to deposit ECM for 6 days. Deposited ECM was decellularized and its fibronectin (FN) content visualized by immunofluorescence **(C)**. FN-positive fibres were quantified using high content microscopy and analysis **(C)**. Values are mean±SEM, statistical significance is indicated by *p* value, data were analyzed by unpaired t-test. Each point on the graph represents the mean relative fold change from an individual experiment. (**D-F**) Control (siNT) and eIF4A2 (si4A2) knockdown fibroblasts were incubated with ‘heavy’ SILAC amino acids (Arg10 & Lys8) for 16h (pulsed-SILAC; **(D)**) or were left unlabeled (label-free;**(E)**). Lysates were digested with trypsin and tryptic peptides analyzed using mass spectrometry to reveal the influence of *Eif4a2* knockdown on the newly synthesized (pulsed-SILAC **(D)**) and total (label-free **(E)**) proteomes. Data are plotted as differences in intensity (abscissa) and statistical significance (ordinate) between *Eif4a2* knockdown and control (si4A2 – siNT). ECM-related (matrisome) components are highlighted by red dots in (**D) & (E**), and the heatmap in (**F**) ranks the intensity differences for the indicated matrisome components between control and *Eif4a2* knockdown cells (si4A2 – siNT) as determined by pulsed-SILAC (upper row in (**F**)) and label-free (lower row in (**F**)) proteomics.

Despite this landscape of increased translation, label-free proteomics indicated that eIF4A2 knockdown did not affect ‘steady-state’ levels of ECM proteins (Fig. 4E,F), suggesting that protein turnover might also be influenced by eIF4A2 levels. We employed a tandem mass tag (TMT)-SILAC approach to quantify protein degradation/turnover on a global scale^28^, which we term ‘turnover-SILAC’. Here, a pulse of labelled amino acids is introduced and the rate at which these are incorporated into newly synthesized proteins (synthesis rate) is compared with the reduction in abundance of the corresponding unlabeled proteins (degradation/turnover rate) both of which may be expressed as half-times (t_1/2_) (Fig. 5A,B). Turnover-SILAC indicated that eIF4A2 knockdown decreased the t_1/2_s of a cohort of proteins, many of which were adhesion- and/or ECM-related components (Fig.5B; Fig. S5). Further analyses revealed that eIF4A2 knockdown led to significantly faster synthesis and degradation rates for this ECM-related cohort of proteins (Fig. 5C). This indicated that the synthesis and degradation rates for many ECM and ECM-related components are closely linked, and that both these rates are coordinately increased following eIF4A2 knockdown.

**Fig. 5.**
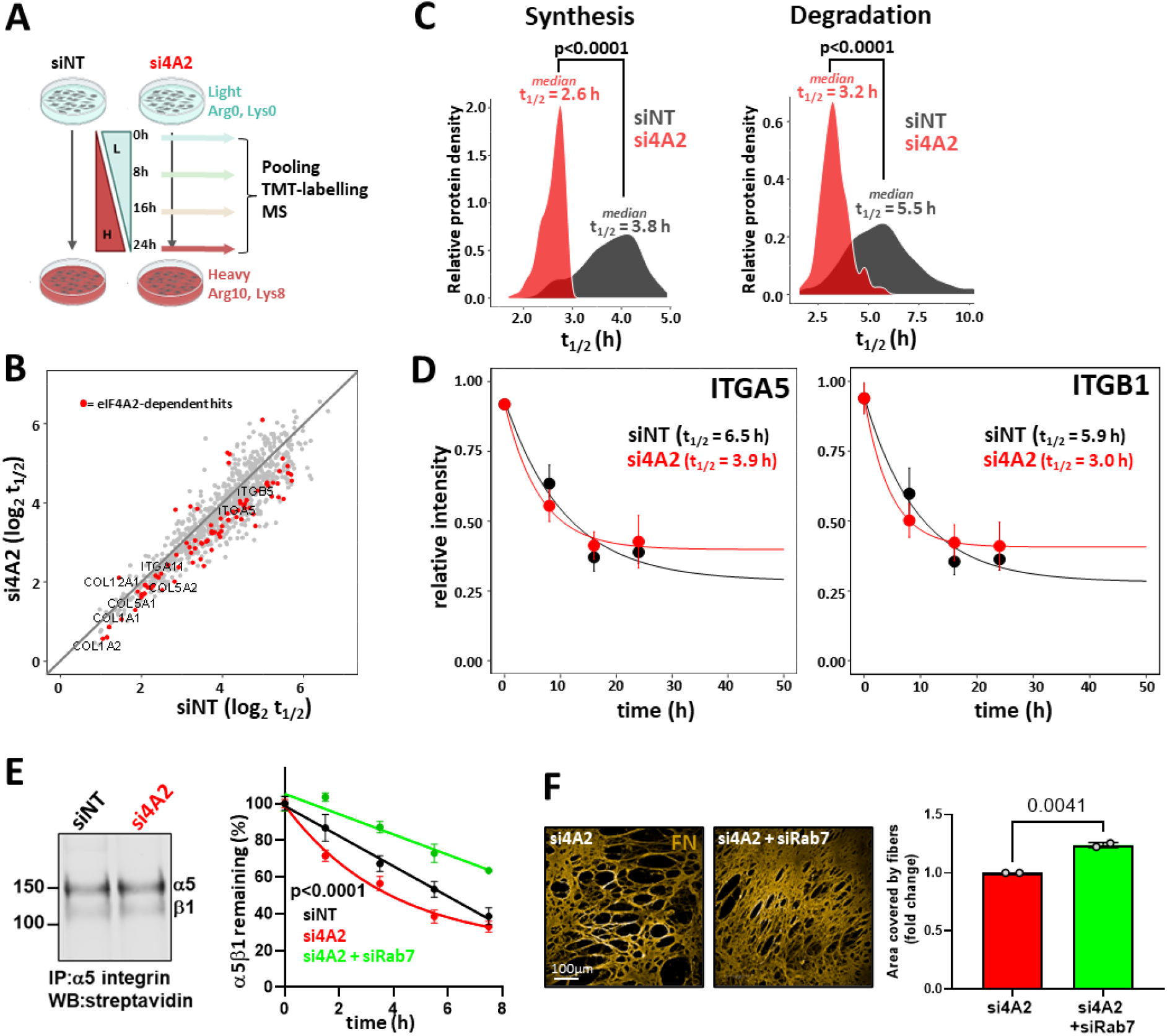
eIF4A2 knockdown promotes increased membrane protein turnover to compromise α5β1 integrin function. (**A-D**) Control (siNT) and *Eif4a2* (si4A2) knockdown fibroblasts were incubated with ‘heavy’ SILAC amino acids for the indicated times (**A**), and tryptic peptides were tandem mass-tagged (TMT) and analysed using mass spectrometry-based proteomics. A protein turnover map (expressed as protein half-life (t1/2)) was derived from the rates at which light-labelled peptides were supplanted by their heavy labelled counterparts (**B**). Proteins with significantly altered t1/2 were ranked (see Fig. S5) and highlighted as red dots on the map (**B**). The synthesis and degradation/turnover rates from the cohort of proteins listed in figure S5 are displayed in (**C**); the respective median t1/2 are indicated and significance value (Wilcoxon test) is displayed. Degradation rates of the constituent subunits of α5β1 integrin – ITGA5 and ITGB1 - are displayed in (**D**). (**E-F**) Senescent fibroblasts in which eIF4A2 had been knocked down either alone (si4A2) or in combination with Rab7 (si4A2 + siRab7) were surface-biotinylated using sulfo-NHS-biotin at 4°C and were either placed at 37°C for the indicated times (**E**) or were allowed to deposit ECM for 6 days (**F**). Levels of biotinylated α5β1 integrin were then determined using immunoprecipitation (IP) followed by western blotting (WB) with HRP-streptavidin (**E**; left panel) or by capture-ELISA (**E**; right graph). WB image is representative of two independent experiments. ELISA data are the mean±SEM of three independent experiments analyzed by nonlinear regression. In (**F**) FN-positive fibres were visualized using immunofluorescence and quantified using high content microscopy and analysis as for Fig. 4C. Each point on the graph represents the mean relative fold change from an individual experiment.

ITGA5 and its partner β1 (ITGB1) integrin are specific examples of ECM-related proteins whose turnover rates (as evidenced by reduced t_1/2_ for degradation) were increased following eIF4A2 knockdown (Fig. 5D). To confirm this, we performed cell-surface labelling/chase followed by capture-ELISA^29^. This indicated that eIF4A2 knockdown significantly increased degradation of surface-labelled α5β1 in senescent cells without altering steady-state levels of the α5β1 heterodimer at the cell surface (Fig. 5E), consistent with the results from our turnover-SILAC proteomics. Taken together, these quantitative proteomic and cell biological analyses indicate that eIF4A2 represses synthesis of a cohort of proteins which includes ECM-related components. Interestingly, when this repression is relieved by *Eif4a2* deletion, cells respond by coordinately increasing degradation/turnover of ECM-related proteins.

We hypothesized that increased degradation of α5β1 integrin might be responsible for reduced FN deposition following *Eif4a2 loss*. α5β1 integrin degradation is controlled by its trafficking through endosomes^29–32^. Therefore, we knocked-down RAB7, a GTPase key to lysosomal routing of active integrins^33^. This reversed the ability of eIF4A2 knockdown to increase α5β1 integrin degradation (Fig. 5E) and restored FN deposition (Fig. 5F). Taken together, these data indicate that translational repression by eIF4A2 enables FN deposition by moderating lysosomal routing and degradation of α5β1 integrin.

### Inhibition of protein synthesis restores ECM deposition and HCC initiation following *Eif4a2* deletion

Pharmacological inhibition of mRNA translation is being considered as an approach to oppose tumour growth. Strategies to target eIF4F show promise in preclinical models of various cancer types and agents targeting protein synthesis, such as mTOR inhibitors, can oppose growth of established tumours in mice and humans^34–36^. However, in view of our finding that *increased* translation rates *reduce* fibronectin deposition and slow HCC progression, we assessed the consequences of reducing protein synthesis on senescence-induced FN deposition (Fig. 6A) and HCC initiation (Fig. 7A. Firstly, we used pSILAC proteomics to confirm that the mTOR inhibitor, rapamycin - an established inhibitor of mRNA translation - reversed the ability of eIF4A2 knockdown to increase ECM-related protein synthesis (Fig. 6B). Daily administration of rapamycin restored the ability of hepatocyte-specific *Mdm2* deletion to drive fibronectin deposition following *Eif4a2* deletion (Fig. 6A), consistent with a causal link between increased mRNA translation and reduced fibrosis *in vivo*. We, therefore, investigated whether administration of rapamycin might similarly influence the FN deposition which precedes establishment of HCC. We induced hepatocyte-specific activation of β-catenin/MYC and deletion of *Eif4a2*. As previously described (Fig. 1), this generated a GS- and p21-postive population of hepatocytes which were unable to increase FN deposition in the liver (Fig. 7A,B) 30 days following AAV-TBG-Cre injection and displayed only limited progression to proliferating HCC at day 90 (Fig. 7C). Daily administration of rapamycin during early tumorigenesis (between 5 and 30 days following AAV-TBG-Cre injection) restored both liver FN deposition (Fig. 7B) and progression to proliferating HCC following *Eif4a2* deletion; the latter evidenced by the abundant presence of tumours in rapamycin-treated *Eif4a2* knockout livers 90 days following AAV-TGB-Cre injection (and 60 days following rapamycin withdrawal) (Fig. 7C).

**Fig. 6.**
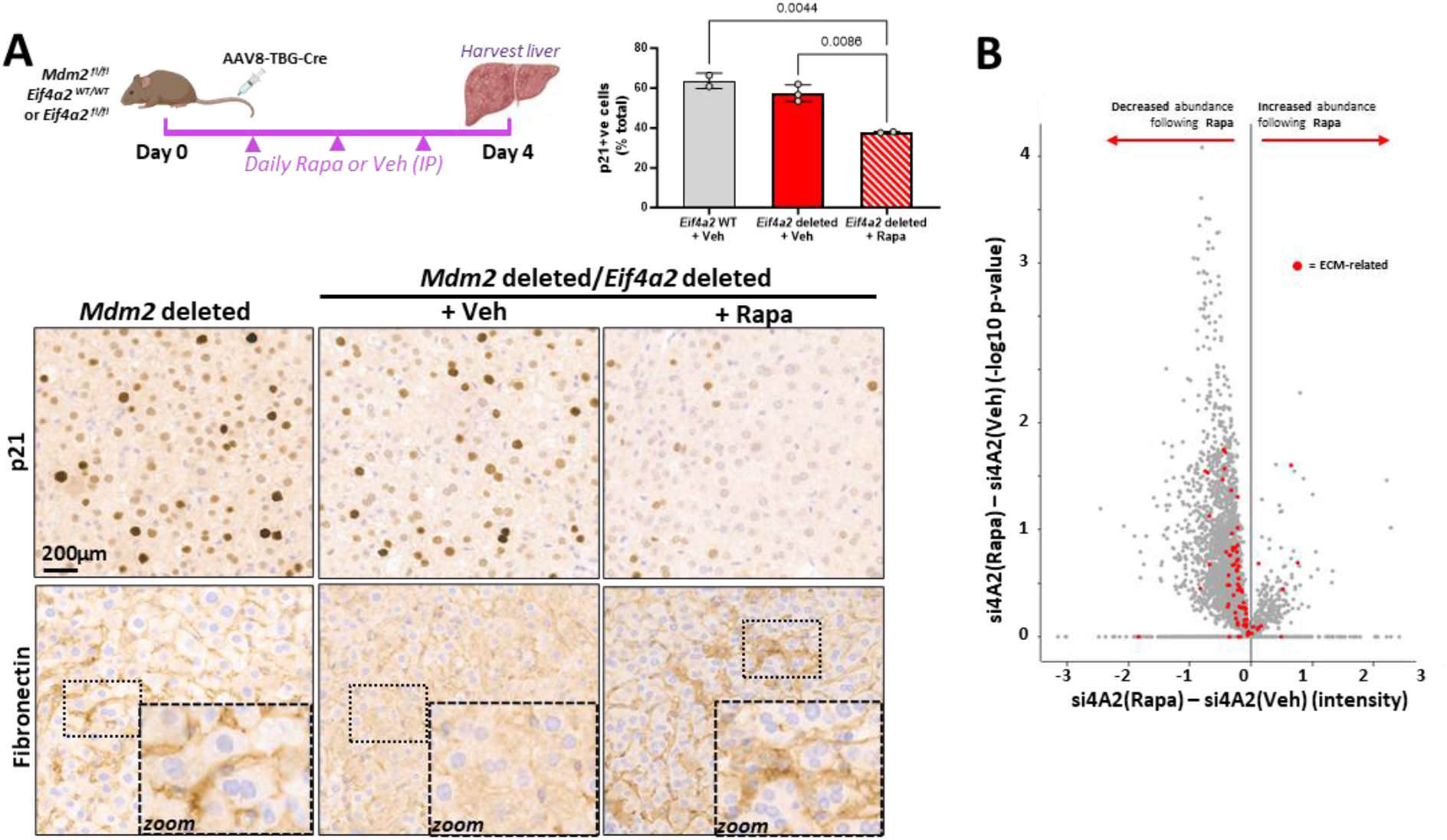
Inhibition of protein synthesis restores fibronectin deposition following *Eif4a2* deletion. **(A)** *Mdm2*^fl/fl^ mice that were either *Eif4a2*^WT/WT^ or *Eif4a2*^fl/fl^ were injected with AAV-TBG-Cre (2 X 10^11^ GC per mouse; a titre sufficient to evoke recombination in >90% hepatocytes). Rapamycin or vehicle control were administered daily by IP injection and mice were sacrificed 4 days later. Fibronectin and p21 were visualized in zone 2 of the liver using immunohistochemistry. Each point on the graph represents data from an individual mouse liver. Data were analyzed by one-way ANOVA. **(B)** eIF4A2 (si4A2) knockdown fibroblasts were incubated with ‘heavy’ SILAC amino acids (Arg10 & Lys8) for 16h in the presence of rapamycin (Rapa) or vehicle control (Veh). Lysates digested with trypsin and tryptic peptides were analyzed using mass spectrometry to reveal the influence of rapamycin treatment on the newly synthesized (pulsed-SILAC) proteome of si4A2 knockdown cells. Data are plotted as differences in intensity (abscissa) and statistical significance (ordinate) between rapamycin treated and vehicle control (si4A2(Rapa) – si4A2(Veh)). ECM-related (matrisome) components are highlighted by red dots.

**Fig. 7.**
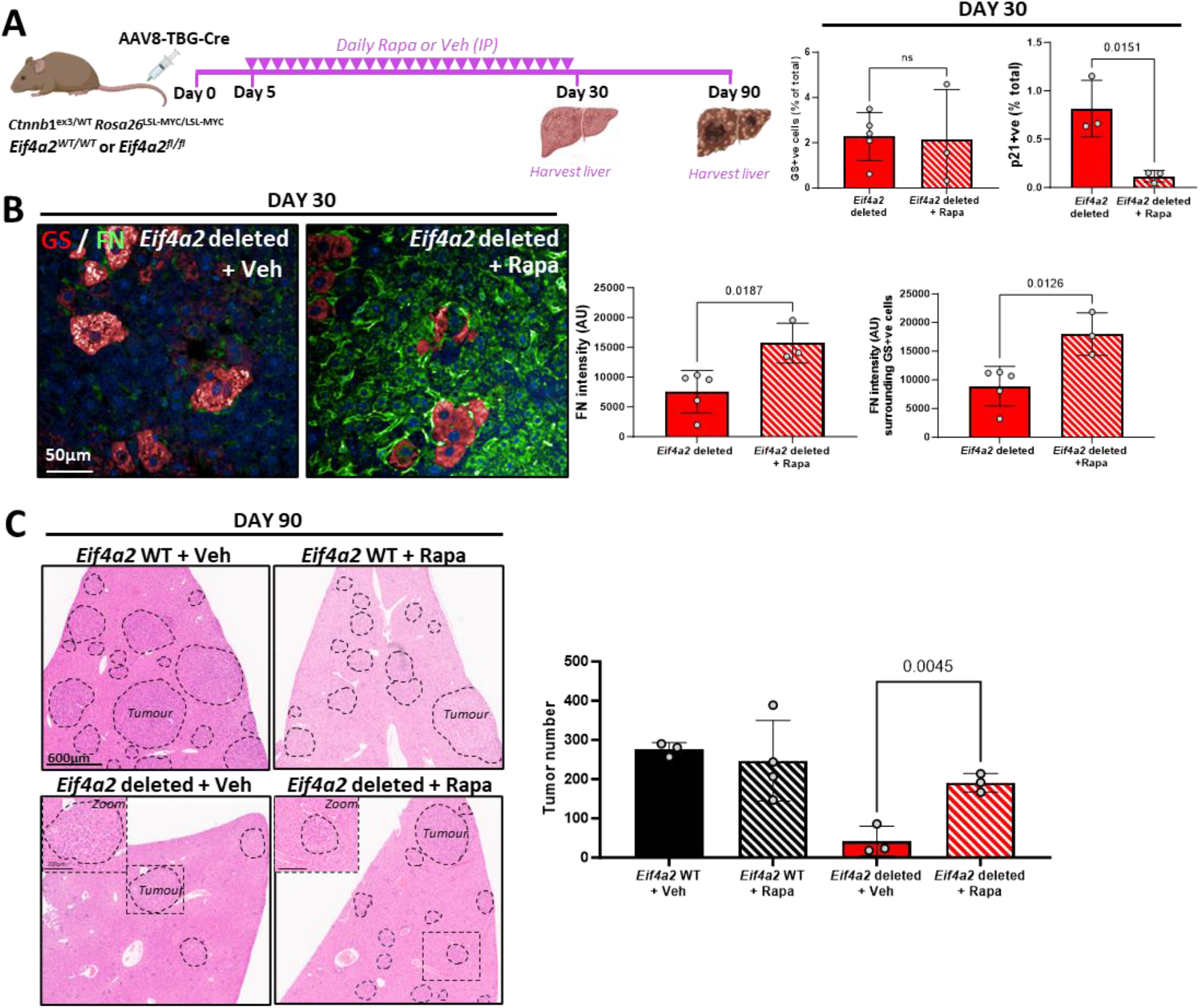
Inhibition of protein synthesis restores HCC initiation following *Eif4a2* deletion. *Ctnnb*1^ex3/WT^; *Rosa26*^LSL-MYC/LSL-MYC^ mice that were either *Eif4a2*^WT/WT^, or *Eif4a2*^fl/fl^ were injected (i.v.) with AAV8-TBG-Cre (6.4 X 10^8^ GC per mouse; a titre sufficient to evoke recombination in ≈5% hepatocytes). 5 days following this, rapamycin or vehicle control were administered daily by IP injection for 25 days (**A**). Mice were sacrificed 30 (**A, B**) or 90 (**C**) days following AAV8-TBG-Cre injection. GS, p21, and fibronectin (FN) were visualized in zone 2 of the liver by immunofluorescence (**A, B**) and the number of tumours determined by manual scoring (**C**). Each point in the graphs represents the quantification from an individual mouse liver, values are mean±SEM, statistical significance is indicated by *p* value, and determined by one-way ANOVA.

## Discussion

Hepatocyte senescence is driven by toxic and genetic insults^21^ and cell cycle exit and the SASP associated with oncogene-induced senescence has likely evolved to perform tumour suppressive roles. Cell cycle exit arrests proliferation of hepatocytes following activation of oncogenic signaling, and the SASP promotes T-cell and monocyte/macrophage ingress to execute clearance of damaged and transformed hepatocytes and limit HCC development^23, 24, 37^. Moreover, it is likely that the SASP contributes to establishment of regenerative niches in the liver to replace hepatocytes following immune clearance, and the presence of FN in the hepatocyte SASP is consistent with this. Thus, eIF4A2 may contribute to liver repair by moderating secretory outputs to maintain integrin function in damaged hepatocytes, thus permitting FN deposition and regenerative niche formation. Whilst regeneration is promoted elsewhere, the damaged hepatocyte may then be cleared by the immune system and the regenerative niche shutdown. We submit that when hepatocyte damage is multifocal, eIF4A2- and α5β1 integrin-dependent processes lead to extensive and persistent deposition of FN which promotes senescence escape and generates regenerative niches to drive HCC progression. Conversely, in tissues where regenerative niches are constitutive and do not need to be damage-activated, such as the gut, there may be less requirement for eIF4A2 in tumour progression. Indeed, eIF4A2 is not required for establishment of adenomas in the gut, but strongly opposes their progression [see accompanying manuscript].

Tissue expression of eIF4A paralogues suggests roles for eIF4A1 and eIF4A2 in supporting proliferative and secretory phenotypes respectively. The switch from proliferative to secretory phenotypes – such as during oncogene-induced senescence - presents challenges to the cell and it is likely that eIF4A2 assists with meeting these demands. A major challenge to the cell is posed by the hugely increased flux of soluble and integral membrane proteins through the ER and Golgi which accompany acquisition of the SASP. Firstly, as the maintenance of correct protein folding and assembly of multi-subunit complexes depends on the dynamics of protein synthesis, the quality of both secreted and membrane components may be compromised if translation initiation and elongation rates at the ER increase in an immoderate manner. Secondly, abnormally large increases in secretory protein production may overload the capacity of the plasma membrane and secretory pathway leading to ‘crowding’ and aberrant function/signaling of plasma membrane receptors. It is likely, therefore, that a key role of the eIF4A2/CCR4-NOT complex is to moderate translation of mRNAs encoding secretory and membrane proteins during acquisition of highly secretory phenotypes to maintain quality control of the secretome and proper homeostasis of the membrane proteome. Clearly, the endosomal pathway has the capacity to compensate for poorly tempered increases in translation because, as we show, cells coordinately upregulate internalization and lysosomal routing of the same cohort of (ECM-related) proteins whose synthesis is overdriven by eIF4A2 depletion. However, although this normalizes levels of these proteins at the plasma membrane – thus avoiding ‘crowding’ – it does not restore integrin function to eIF4A2 knockdown cells and would be unlikely to correct errors in protein folding/subunit assembly.

So, how could eIF4A2/CCR4-NOT moderate translation in a way that enables proper folding/function of membrane proteins^38^. The CCR4-NOT complex assists in targeting mRNAs to the ER, but also to sensing codon optimality^39^ and this may be why it slows mRNA translation. Therefore, CCR4-NOT may enable translating ribosomes to sense when they have reached a suboptimal codon and to pause for sufficient time to allow the nascent polypeptide chain to fold before translation proceeds^40^. The addition of rapamycin can restore integrin function following *Eif4a2* deletion and we believe this is consistent with a mechanism in which the role of the CCR4-NOT/eIF4A2 complex is to slow secretory protein synthesis to allow their proper folding, assembly, processing, and function whilst cells acquire highly secretory phenotypes. Finally, we believe our study highlights an emerging paradigm outlining a mechanistic relationship between a tissue’s ability to support high secretory function and its propensity to foster malignancies. Indeed, it is the hepatocyte’s capacity to engage eIF4A2 and, thereby, ameliorate cellular stress associated with production of the SASP that allows hepatocytes which have entered oncogene-induced senescence to reenter the cell cycle and form aggressive carcinoma.

This study demonstrates that pharmacological approaches to reduce protein synthesis during early tumorigenesis can increase establishment of FN-rich tumour initiation niches leading to progression from a pre-malignant OIS state to proliferating HCC. However, we do not believe that this model precludes a role for mRNA translation in driving tumour size. Indeed, although rapamycin treatment can restore tumour initiation following *Eif4a2* deletion, the resulting tumours are smaller than those from untreated mice (Fig. 7E). Thus, although inhibition of protein synthesis may be an effective way to reduce tumour biomass and the growth of established tumours^34–36^, it is important to consider that high levels of translation can extend a period of senescence occurring following oncogene activation. Thus, the use of drugs which reduce mRNA translation, if administered during this period of oncogene-induced senescence, may awaken senescent cells and promote tumour progression.

## Author contributions

Experimental design: MM, LP, LM, TS, SM, TGB, MB, JCN

Methodology: MM, LP, LM, TS, SM, MM, RCLS, DS, SB, KH, SL, SZ, LM, SG, JW, RP, CN,

Investigation: MM, LP, LM, TS, SM, MM, RCLS, DS, AW, SB, KH, SL, LM, SG, JW, RP, GK, CN, JCN

Visualization: MM, LP, LM, CN

Funding acquisition: OJS, TGB, MB, JCN Writing – original draft: MM, LP, JCN

Writing – review & editing: MM, LP, LM, JW, OJS, TGB, MB, JCN

## Data and materials availability

Links to relevant polysome profiling and proteomic data are provided in the supplementary text (material and methods). All other primary data, reagents, and mice are available from JN upon request.

## Materials and Methods

The data that support the findings of this study are available on request from the corresponding author, [JN]. The mass spectrometry proteomics data have been deposited to the ProteomeXchange Consortium via the PRIDE partner repository with the dataset identifiers: PXD044626 (Pulsed-SILAC); PXD044651 (label-free) and PXD044585 (TMT-SILAC) The RNAseq data from Polysome profiling experiment has been deposited at GEO accession GSE243755. To access these data, please go to https://www.ncbi.nlm.nih.gov/geo/query/acc.cgi?acc=GSE243755 and enter token shqjioowbhkddgl into the box.

### Mice and treatments

All animal experiments were performed in accordance with the UK Animal (Scientific Procedures) Act 1986 and EU direction 2010 and the UK Home Office licences 70/8891 and PP0604995. All procedures were ethically reviewed by the animal welfare and ethical review board, University of Glasgow. Animals were housed under controlled conditions (19-22°C room temperature, 45-65% humidity, pathogen free, 12h light/dark cycle) with access to food and water *ad libitum*. All animals received environmental enrichments including gnawing sticks, plastic tunnels, and nesting material. To reduce pain, suffering and distress, single use needles were used when administering virus and drug treatments, and non-adverse handling techniques were adhered to. All mice used were bred in-house and were of a mixed background. Genotyping was performed by Transnetyx Inc. using ear notches taken for identification purposes at weaning (approximately 3 weeks of age). Targeting constructs for the conditional *Eif4a1* and *Eif4a2* alleles were generated used to generate the *Eif4a1* and *Eif4A2* floxed mice as described in the accompanying manuscript.

The mice used for experimental purposes were genetically modified with the following alleles: *Ctnnb1*^tm1Mmt^ (*Ctnnb1*^ex3^), Gt(*Rosa*)26S or tm1(MYC)Djmy (R26^LSL-MYC/LSL-MYC^), *Mdm2*^flox/flox 41^, Itga5^flox/flox 20^, *Eif4a1*^flox/flox^ and *Eif4a2*^flox/flox^ (generated in-house as described in the accompanying manuscript). Mice were induced between 8 and 12 weeks of age. Males weighed a minimum of 20g and females, a minimum of 18g, at the time of viral induction. Animal technicians administering the virus and scientists monitoring for clinical signs were blinded to the genotype of the mice. Mice were sampled at specific time points (see figure legends for details) or at a defined clinical endpoint. Mice that died due to tumour hemorrhaging were censored from survival cohorts and any downstream analyses. Unless otherwise stated, liver tissue from the median lobe was sampled in neutral buffered saline containing 10% formaldehyde and liver tissue from the left lobe was snap frozen on dry ice. The number of biological replicates was ≥ 2 mice per cohort for all experiments (see Figure legends for details). For the HCC model, male mice were injected with 6.4 X 10^8^ GC/mouse of AAV8.TBG.PI.Cre.rBG (AAV-TBG-Cre, Addgene, 107787-AAV8) diluted in 100μL PBS via the tail vein. The Log-rank Mantel-Cox test was used to determine statistical significance. For the hepatic senescence model: female mice were injected with 2 X 10^11^ GC/mouse of AAV8.TBG.PI.Cre.rBG (AAV-TBG-Cre) diluted in 100μL PBS via the tail vein and tissues were harvested at 4 days post viral induction. For rapamycin drug treatments, mice were administered rapamycin (10mg/kg) diluted in PBS with 5% ethanol, 5% PEG400 (Sigma) 5% Tween 20 (Sigma) or vehicle control via daily IP injection and tissues were harvested at specific end points (see figure legends for details).

### Cell lines, antibodies and reagents

Human lung fibroblasts, IMR90-ER:RAS cells (IMR90s) expressing a chimeric fusion protein that activates upon treatment with 4-hydroxytamoxifen (4OHT) were a kind gift from Juan Carlos Acosta, University of Cantabria, Spain, and are described in^42^. IMR90s were cultured in Dulbecco’s modified Eagle medium (DMEM, Sigma) without phenol red and supplemented with 10% fetal bovine serum (FBS), 2 mM glutamine, 100 IU/ml penicillin and 100 μg/ml streptomycin. Oncogene-induced senescence (OIS) was induced using 100nM 4OHT (Sigma) for 72h. Immortalized human breast fibroblasts (iCAF, ^43^) were used for mass spectrometry-based proteomics. Cells were cultured in DMEM supplemented with 10% fetal bovine serum (FBS), 2 mM glutamine, 100 IU/ml penicillin and 100 μg/ml streptomycin. All cell lines were tested for the presence of mycoplasma. For Western blots, the following antibodies were used at a dilution of 1:1000 (unless otherwise stated): eIF4A1 (abcam 31217), eIF4A2 (abcam 31218), ITGA5 (BD Biosciences 610633), GAPDH 1:5000 (Sigma G8795) and Vinculin (Millipore MAB3574-C).

### siRNAs and transfections

IMR90 cells were seeded at 6000 cells/cm^2^ and treated with 100nM 4OHT for 72h. Cells were forward transfected with 30nM siRNA using Lipofectamine 2000 (Thermo Fisher) according to the manufacturer’s instructions and downstream experiments were performed 72h post transfection. CAF siRNA transfections were performed by nucleofection using the Amaxa kit R (Lonza) and downstream experiments were performed 5 days post transfection. SMARTPool siRNA oligos targeting eIF4A2, eIF4A1 and Rab7 were purchased from Dharmacon.

### Cell-derived matrix generation & fibronectin analysis

Extracellular matrix (ECM) from senescent IMR90 cells was generated as previously described^44^. Briefly, cells were treated with 100nM 4OHT for 72h and then transfected with siRNAs (as described above). 24h post transfection cells were re-plated at 100% confluency in gelatin coated 6-well glass bottom plates (MatTek) and treated with ascorbic acid every two/three days to promote matrix deposition. At 8 days post-transfection, cells were removed using denudation buffer (20mM NH_4_OH, 0.5% Triton X-100 in PBS) and the ECM was fixed using 4% paraformaldehyde (MP biomedicals). For fibronectin analysis, cell-derived matrices were blocked with 1% BSA and stained with FN antibody (BD Pharmingen 610078 1:100) for 2h at RT and then with secondary antibody, Alexa-555 (Thermo Fisher 1:400) for 45 min at RT. Fibronectin was imaged at ×20 magnification using the Opera Phenix High-Content Screening System (Harmony High-content Imaging and analysis software version 4.9, PerkinElmer) and fibronectin fibers were quantified using Columbus Image Data Storage and Analysis System (PerkinElmer version 2.8.0).

### ELISA-based degradation assay

Senescent IMR90 cells were transfected with 30nM siRNA, as described above. 72h post transfection cells were surface labelled at 4°C with 0.13 mg/ml NHS-SS-biotin (Pierce) and then warmed to 37°C. The proportion of α5β1 remaining was determined by capture-ELISA using Maxisorp (Nunc) plates coated with antibodies recognizing each receptor, as described previously^29^. The following antibodies were used at a concentration of 5µg/ml in 0.05M Na_2_CO_3_ pH 9.6 at 4°C: α5 integrin clone VC5 (Pharmingen) for total α5β1 (human) and antiCD71 (Pharmingen) for TfnR.

### Tissue Immunostaining and Microscopy

#### Immunohistochemistry (IHC)

All Haematoxylin & Eosin (H&E) and immunohistochemistry (IHC) staining was performed on 4µm formalin fixed paraffin embedded sections (FFPE) which had previously been heated at 60⁰C for 2 hours. The following antibodies were used on an Agilent AutostainerLink48, GS (HPA007316, Sigma), eIF4A1 (ab31217, Abcam), and eIF4A2 (ab31218, Abcam). Sections were loaded into an Agilent pre-treatment module to be dewaxed and undergo heat induced epitope retrieval (HIER) using high pH target retrieval solution (High pH TRS K8004, Agilent) (for eIF4A1 and eIF4A2) or low pH target retrieval solution (Low pH TRS K8005, Agilent) (for GS). The sections underwent peroxidase blocking (S2023, Agilent) for 5 minutes and washed with flex buffer. Primary antibody was applied for 35 mins and secondary antibody (Rabbit Envision K4003, Agilent) for 30 minutes. Sections were then incubated with liquid DAB (K3468, Agilent) for 10 minutes and counterstained with haematoxylin z (RBA-420100A, CellPath). α-Smooth muscle actin (αSMA) (19245, Cell Signaling), p21 (ab107099, Abcam), Ki67 (12202S, Cell Signalling) and Fibronectin (ab2413, Abcam) antibodies were stained on a Leica Bond Rx. Sections for p21, Ki67 and Fibronectin underwent antigen retrieval using ER2 solution (AR9640, Leica) for 20 minutes at 100°C, α-SMA for 30 minutes. Sections were rinsed with Leica wash buffer (AR9590, Leica) before peroxidase block was performed using an Intense R kit (DS9263, Leica). Sections for p21 had blocking solution applied from the Rat ImmPRESS kit (MP-7404, Vector Labs) for 20 minutes. Primary antibody was applied for 35 minutes, and secondary antibody applied for 30 minutes. αSMA, Ki67 and Fibronectin had Rabbit EnVision applied. Sections for p21 had Rat ImmPRESS secondary solution applied. Sections were visualized using DAB and then counterstained with haematoxylin in the Intense R kit. H&E staining was performed on a Leica autostainer (ST5020) with Haem Z (RBA-4201-00A, CellPath) and Putt’s Eosin (in-house). To complete H&E and IHC staining sections were rinsed in tap water, dehydrated through graded ethanol’s and placed in xylene. The stained sections were mounted onto coverslips in xylene using DPX mountant (SEA-1300-00A, CellPath).

#### Immunofluorescence staining

antigen retrieval was performed using Citrate Buffer (10mM Citric Acid, pH 6, 0.05% Tween) for 20 minutes at 90°C and tissue was blocked in 1% BSA for 1h at RT and incubated with primary antibody overnight at 4°C. Tissue was incubated with secondary antibodies (Alexa-488 or Alexa-647, Thermo Fisher, 1:400) before being mounted with VECTASHIELD containing DAPI (Vector Laboratories). Antibodies used were: eIF4A2 (abcam 31218, 1:100), GS (BD Biosciences 610517, 1:1000) and ITGA5 (Merck AB1928, 1:1000). Fibronectin (abcam 2413, 1:100) and p21 (abcam 107099, 1:100) were detected using a Tyramide Signal Amplification (TSA-FITC) system (Perkin Elmer, NEL741B001KT, 1:50).

#### High-content automated microscopy

Stained slides were imaged at ×20 magnification using the Opera Phenix High-Content Screening System (Harmony High-content Imaging and analysis software version 4.9, PerkinElmer) and images were quantified using Columbus Image Data Storage and Analysis System (PerkinElmer version 2.8.0). Fibronectin intensity surrounding GS+ve cells refers to fibronectin deposited 2-7μm from the cell membrane of GS+ve cells.

#### IHC scanning and analysis

Stained slides were scanned using the Leica Aperio AT2 slide scanner and analyzed using HALO® image analysis software. Tumour burden was quantified in H&E-stained sections using HALO® Area quantification v2.4.2. Alpha-SMA and p21 protein expression levels were quantified using HALO® cytonuclear v2.0.9 analysis trained to quantify the optical density of cytosolic (α-SMA) or nuclear (p21) staining.

### Statistical analysis

All statistical analysis was performed using GraphPad Prism V9.5.1 software (see figure legends for details). In all instances, *P* values less than or equal to 0.05 were considered significant. Unless otherwise stated, a T-test or One-way-ANOVA was used, depending on the number of comparisons. The degradation assay was analyzed using a non-linear regression and adjusting the one-phase decay curve to the data. Error bars represent mean ± standard error of the mean (SEM).

### Polysome profiling & sequencing

Mice were sacrificed by CO_2_ following with cervical dislocation and whole livers were harvested and ground in a liquid nitrogen chilled Cryocooler (OPS diagnostics). Sample powder was then lysed for 5 minutes on ice using lysis buffer (15 mM TrisHCl (pH 7.4), 15 mM MgCl_2_, 0.15 M NaCl, 1% Triton X-100, 0.05% Tween-20, 0.5mM DTT, 5mM NaF, 2%DDM, 2μg/ml TurboDNAse (Thermo Fisher), 200U/ml Ribolock, 0.1 mg/ml cycloheximide and protease inhibitors) and then spun at 13000rpm for 5 minutes at 4°C. 40μl of lysate were separated for total RNA sequencing, added to 500µL of Trizol (Thermo Fisher) and RNA was extracted following manufacturer instructions. Then, 300 μl lysate was loaded onto a 10–50% sucrose gradient and spun at 38,000 rpm for 2 h at 4 °C. Gradient fractions were collected into 3 ml 7.7 M GuHCl, 8 µl glycogen, and 4 ml 100% ethanol added and then precipitated for > 24h at – 20 °C. RNA was pelleted at 4000 rpm for 1 h at 4 °C. The supernatant was removed, and the pellets resuspended in 400 μl RNase-free water. Then, 2 μl glycogen, 40 μl 3 M NaOAc pH 5.2, and 1 ml 100% ethanol was added to the samples, and these were precipitated overnight at – 20 °C. Samples were then pelleted at 13,000 rpm 4 °C for 40 min, washed with 500 μl 75% ethanol, air dried, and resuspended in 30 μl RNase-free water. To check the RNA integrity, equal volumes (3 μl) of each fraction along the gradient were ran on a 1% agarose gel. RNA concentration was quantified using a NanoDrop 2000c Spectrophotometer (ThermoScientific) and subpolysomal and polysomal fractions were pooled. One microgram of RNA of total, subpolysomal, and polysomal RNA were used for sequencing. RNA libraries were prepared using the Corall Total RNA-Seq Library Prep Kit (Lexogen Cat. 095.96) with the Poly(A) Selection Kit (Lexogen Cat. No. 039). Libraries were sequenced on an Illumina NextSeq using the High Output 75 cycles kit (1 x 75 cycles, single index). RNA-seq data were analyzed using the https://github.com/Bushell-lab/RNA-seq pipeline. Briefly, cutadapt was used to remove adaptors from the FASTQ files and UMI tools was used for deduplication of PCR duplicates using the 12nt UMIs at the start of all sequencing reads. Bowtie2 was used to align to the mouse transcriptome and RSEM used for quantification and DESeq2 for differential expression analysis. Gene Set Enrichment Analysis (GSEA) was carried out using R package fgsea version 1.8.0, and comparison gene sets obtained from available sources (Gene Ontology Consortium).

### Proteomic analyses

In all proteomic experiments proteins were reduced with 10mM DTT and subsequently alkylated in the dark with 55 mM Iodoacetamide, both reactions were carried out at room temperature. Alkylated proteins were then cleaved using a two-step digestion: first with Endoproteinase Lys-C (ratio 1:33 enzyme: lysate, Promega) for 1 hour at 37 °C then with Trypsin (ratio 1:33 enzyme: lysate, Promega) overnight at 37 °C. Label-free samples were desalted after digestion using StageTip^45^ prior to MS analysis. For pulsed Stable Isotope Labelled Amino Acid in Culture (p-SILAC) of CAFs treated with Rapamycin, 100 µg of lysate were digested and fractionated into six fractions using an Agilent AssayMAP Bravo equipped with Reversed Phase RP-S cartridge (Agilent). The “Protein Sample Prep WorkBench Fractionation application” was used with default settings. Cartridges were initially primed with 90 % acetonitrile (ACN) and equilibrated with 20mM ammonium formate. Digested proteins were then loaded and eluted into six fractions with buffers containing increasing ACN percentages (15, 20, 25, 40 and 80%), all buffers were adjusted to pH 10 using ammonium formate. The p-SILAC experiment for the control and eIF4A2 knockdown fibroblasts, was carried out by digesting 300 µg of lysate, and resulting tryptic peptides were directly fractionated using offline high pH reverse phase chromatography. For the p-SILAC-TMT samples, digested peptides were differentially labelled using Tandem Mass Tag (TMT) 16-plex reagent kit (Thermo Scientific) following manufacturer protocol. The reaction was carried out at room temperature for 2 hours and quenched with 5% hydroxylamine. Fully labelled peptide samples were mixed in equal amount and desalted using Sep Pak C18 solid-phase extraction cartridges (Waters) prior to offline high pH reverse phase chromatography.

#### Offline HPLC fractionation (p-SILAC and p-SILAC-TMT samples)

Digested peptides from pSILAC and p-SILAC-TMT experiments were fractionated using high pH reverse phase chromatography. The separation was carried out on a C18 column (150 × 2.1 mm – Kinetex EVO (5 μm, 100 Å)) using an Agilent, LC 1260 Infinity II HPLC system. Elution solvents were A (98% water, 2% ACN) and B (90% ACN, 10% water), both adjusted to pH 10 using ammonium hydroxide. Peptides were separated over a two-step gradient at a flow rate of 200 µL/min for a total run time of 76 mins. Column eluate was collected into 21 fractions, each analyzed by mass spectrometry (MS).

#### MS Analysis

Peptides resulting from all digestions were separated by nanoscale C18 reversed-phase liquid chromatography using an EASY-nLC II 1200 (Thermo Scientific) coupled to an Orbitrap Q-Exactive HF (Thermo Scientific) for the p-SILAC of Control and eIF4A2 knockdown fibroblasts samples, or to an Orbitrap Fusion Lumos (Thermo Scientific) for all the other samples. All acquisitions were carried out in data-dependent acquisition mode (DDA) using Xcalibur software (Thermo Scientific). For both mass spectrometers a nanoelectrospray ion source (Sonation) was used for ionization in positive mode. Chromatography was carried out at a flow rate of 300 nl/min using 20 cm (p-SILAC of Control and eIF4A2 knockdown fibroblasts) or 50 cm (for all other samples) fused silica emitters (CoAnn Technologies). All emitters were packed in house with reverse phase Reprosil Pur Basic 1.9 µm (Dr. Maisch GmbH). An Active Background Ion Reduction Device (ABIRD) was used to decrease air contaminants signal level.

#### MS conditions for Label-free samples

Peptides were eluted with a two-step gradient method, over a total run time of 275 mins. Advanced Peak Determination was turned on and Monoisotopic Precursor Selection was set to “Peptide” mode. A full scan was acquired at a resolution of 240000 at 200 m/z, over mass range of 375-1500 m/z. Ions were selected during a 3 sec cycle time using the quadrupole, fragmented in the ion routing multipole, and finally analyzed in the Ion trap using a maximum injection time of 10 ms or a target value of 3e4 ions. Former target ions selected for MS/MS were dynamically excluded for 20 s.

#### MS conditions for p-SILAC samples

Peptides from CAFs treated with Rapamycin fractionated with AssayMAP Bravo, were eluted over optimized gradients for each of the six fractions, for a total run time of 135 minutes per fraction. A full scan was acquired at a resolution of 120000 at 200 m/z, over mass range of 350-1400 m/z. Ions were selected during a 3 sec cycle time using the quadrupole through a 0.7Da window, fragmented in the ion routing multipole, and finally analyzed in the ion-trap. Ions were isolated for fragmentation with a target of 2E4 ions, for a maximum of 20 ms. Ions that have already been selected for MS2 were dynamically excluded for 30 sec. Fractions from offline high pH reverse phase chromatography of p-SILAC experiment to compare control and eIF4A2 knockdown fibroblasts were analyzed using three different gradients optimized for fractions 1-7, 8-15 and 16-21 as described previously^46^ or a total run time of 53 minutes per fraction. A full scan was acquired at a resolution of 60000 at 200 m/z, over mass range of 375-1400 m/z. HCD fragmentation was triggered for the top 15 most intense ions detected in the full scan. Ions were isolated for fragmentation with a target of 5E4 ions, for a maximum of 50 ms, at a resolution of 15,000 at 200 m/z. Ions that have already been selected for MS2 were dynamically excluded for 25 sec.

#### MS conditions for p-SILAC-TMT samples

Fractionated peptides were eluted over three twostep gradients optimized for fractions 1-7, 8-15 and 16-21 as described previously^47^ for a total run time of 185 minutes per fraction. A full scan was acquired at a resolution of 60000 at 200 m/z, over mass range of 350-1400 m/z. Ions were selected during a 3 sec cycle time using the quadrupole through a 0.8Da window, fragmented in the ion routing multipole, and analyzed in the Orbitrap at 50000 resolution at 200 m/z, using a maximum injection time of 120 ms or a target value of 1e5 ions. Former target ions selected for MS/MS were dynamically excluded for 60 s.

#### MS proteomic data analysis

All acquired MS raw data were processed using MaxQuant^48^ version 1.6.14.0 and searched with Andromeda search engine^49^ against the Uniprot *Homo sapiens* database (20,043 entries, 2022). First and main searches were carried out with a precursor mass tolerance of 20 ppm for the first search, 4.5 ppm for the main search and for the MS/MS the mass tolerance was set to 20 ppm. Minimum peptide length was 7 amino acids and trypsin cleavage was selected allowing up to 2 missed cleavage sites. Methionine oxidation and N-terminal acetylation were selected as variable modifications and Carbamidomethylation as a fixed modification. For quantitation of the p-SILAC experiments in MaxQuant, the multiplicity was set to 2 and Arg0/Arg10, Lys0/Lys8 were used for ratio measurement of SILAC labelled peptides. The Arg10 and Lys8 SILAC modifications were specified as variable during processing of p-SILAC-TMT samples. MaxQuant was set to quantify on “Reporter ion MS2”, and TMT16plex was selected as Isobaric label. Interference between TMT channels were adjusted by MaxQuant using the correction factors provided by the reagent manufacturer. The “Filter by PIF” option was activated and a “Reporter ion tolerance” of 0.003 Da was used. False discovery rate was set to 1% for peptides and proteins.

All MaxQuant outputs were processed with Perseus^50^ version 1.6.15.0. The Reverse, Contaminant and “Only identified by site” hits were removed, and only protein groups identified with at least one unique peptide were allowed in all lists of identified protein groups. The protein abundance for label-free samples was measured using label-free quantification algorithm available in MaxQuant^51^ and reported in the ProteinGroups.txt file. Only proteins robustly quantified in all three replicates in at least one group, were included in the list of quantified proteins. For the p-SILAC TMT experiment, an R script (https://github.com/KHodgeSS0/pSILAC-TMT-Rebuild-Script) described previously^52^ was used to calculate the intensities of heavy and light protein using the TMT reporter ion intensities from labelled SILAC reported in the Evidence.txt file. Protein TMT reporter ion intensities were normalized across multiple TMT 16 plex datasets using LIMMA^53^ plugin in Perseus.

#### Analysis of turnover, synthesis and degradation data from TMT-turnover-SILAC

Further data processing and analyses were performed as described previously^28, 52^. Briefly, for turnover analysis, a custom R-script (available on request) was used to convert the per-gene intensity data into fraction ‘heavy’ (F_H_) by dividing TMT-intensities from ‘heavy-labelled’ by the sum of light- and heavy-labelled TMT intensities (F_H_ = I_Lys8,Arg10_ / (I_Lys8,Arg10_ + I_Lys0,Arg0_). For individual synthesis (heavy TMT labels I_Lys8,Arg10_) and degradation (light TMT labels I_Lys0,Arg0_) analysis, intensity data were normalized to timepoints 24 h and 0 h respectively. All data sets were filtered for steadiness i.e., turnover and synthesis data showing increasing trend over time and degradation decreasing trend over time. To calculate the apparent turnover, synthesis and degradation rates *k*, a single exponential equation for labelling kinetics was fitted to the normalized data I_N_, (I_N_ = I_0_ + amplitude * e^-^*^k^* ^* t^). To remove poor quality fits, data for the rate *k* were filtered using a p-value cut-off of 0.1 (two-sided t-test, null-hypothesis of *k* = 0). Next, log2-fold changes of the half-lives were calculated (si4A2/siNT) using standard procedures. Proteins were classified as eIF4A2-dependent or -independent if their p-value was smaller than 0.05 or larger than 0.7 respectively. This resulted in 223 eIF4A2-dependent and 159 eIF4A2 independent genes. Gene set (cellular component) enrichment analysis of eIF4A2-dependent genes was performed using enrichR^54, 55^. In Fig 3 only terms with an FDR < 0.1 are shown.

---

**Fig. S1.**
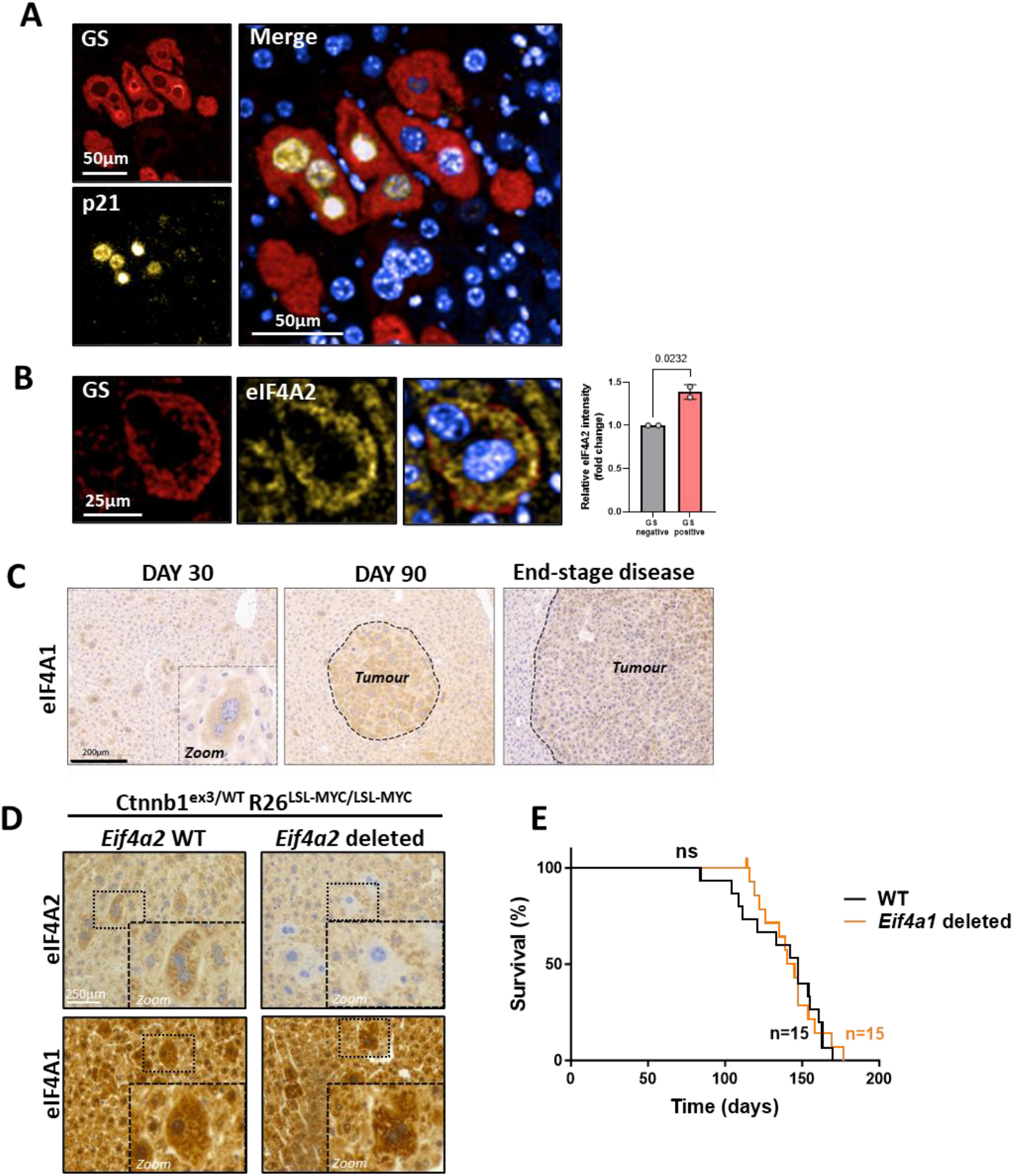
(A-B) Immunofluorescence visualization of p21 and eIF4A2 in HCC initiating cells. *Ctnnb*1^ex3/WT^; *Rosa26*^LSL-MYC/LSL-MYC^ mice were injected (i.v.) with AAV-TBG-Cre (6.4 X 10^8^ genome copies (GC) per mouse; a titre sufficient to evoke recombination in ≈5% hepatocytes). Mice were sacrificed 30 days following this and glutamine synthetase (GS; an indicator of activated β-catenin signaling) was visualized with respect to p21 (A) or eIF4A2 (B) in zone 2 of the liver by immunofluorescence. The intensity of staining for eIF4A2 in GS-positive and GS-negative hepatocytes was determined using high-content analysis - each data point in (B) represents quantification from an individual mouse, mean±SEM, statistical test was student’s unpaired t-test. **(C) Expression of eIF4A1 during HCC progression.** *Ctnnb*1^ex3/WT^; *Rosa26*^LSL-MYC/LSL-MYC^ mice were injected with AAV-TBG-Cre as for A and sacrificed 30 or 90 days following this or were left till tumour endpoint (end-stage disease) was reached. eIF4A1 was visualized using immunohistochemistry. **(D) Knockout of *Eif4a2* in HCC tumor initiating cells.** *Ctnnb*1^ex3/WT^; *Rosa26*^LSL-MYC/LSLMYC^ mice that were either *Eif4a2*^WT/WT^ or *Eif4a2*^fl/fl^ were injected (i.v.) with AAV-TBG-Cre (6.4 X 10^8^ GC per mouse). Mice were sacrificed 30 days following this and eIF4A1 and eIF4A2 were visualized in zone 2 of the liver by immunohistochemistry. **(E) Hepatocyte-specific knockout of *Eif4a1*does not influence initiation or progression of HCC.** *Ctnnb*1^ex3/WT^; *Rosa26*^LSL-MYC/LSL-MYC^ mice that were either *Eif4a1/Eif4a2*^WT/WT^ or *Eif4a1*^fl/fl^ were injected (i.v.) with AAV-TBG-Cre (6.4 X 10^8^ GC per mouse). Mice were sacrificed at tumour endpoint and the time (in days) to endpoint was recorded. Statistical test, Log rank (Mantel-Cox).

**Fig. S2.**
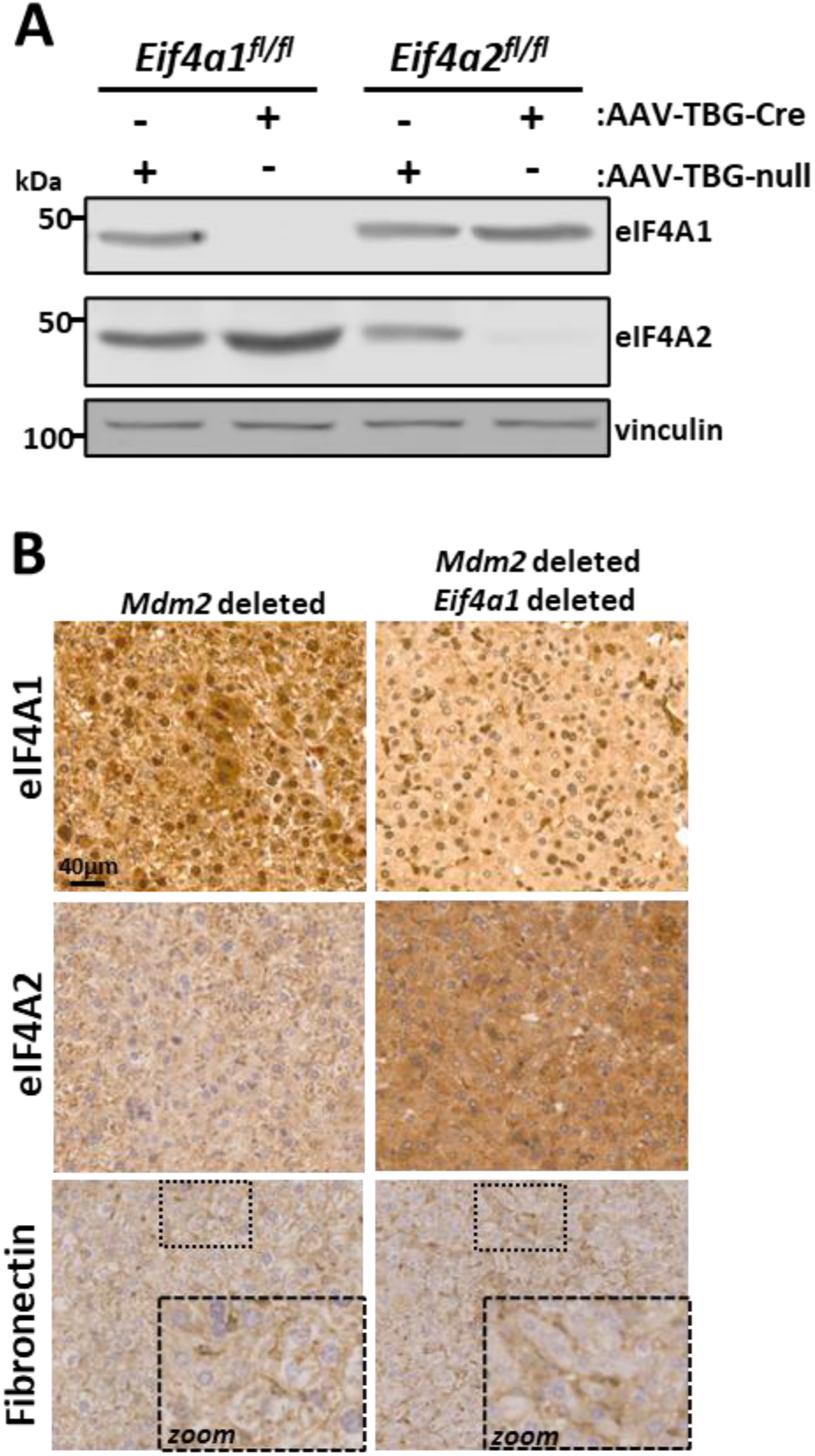
Roles and expression of eIF4As and fibronectin in liver senescence. **(A) Hepatocyte-specific deletion of eIF4A paralogues.** Mice bearing floxed alleles of *Eif4a1* (*Eif4a1*fl/fl) or *Eif4a2* (*Eif4a2*fl/fl) were injected with AAV-TBG-Cre (2 X 1011 GC per mouse; a titre sufficient to evoke recombination in >90% hepatocytes) and sacrificed 30 days later. Livers were removed and homogenized. Levels of eIF4A1, eIF4A2 in liver homogenates were determined by Western blotting with vinculin as loading control. **(B) eIF4A1 deletion does not oppose FN deposition following induction of hepatocyte senescence.** Mice bearing floxed alleles of *Mdm2* (*Mdm2*fl/fl) that were either *Eif4a1* WT (*Eif4a1*WT/WT) or *Eif4a1* floxed (*Eif4a1*fl/fl) were injected with AAV-TBG-Cre as for (**A**). 4 days following this, livers were fixed as eIF4A1, eIF4A2 and fibronectin visualized using immunohistochemistry.

**Fig. S3.**
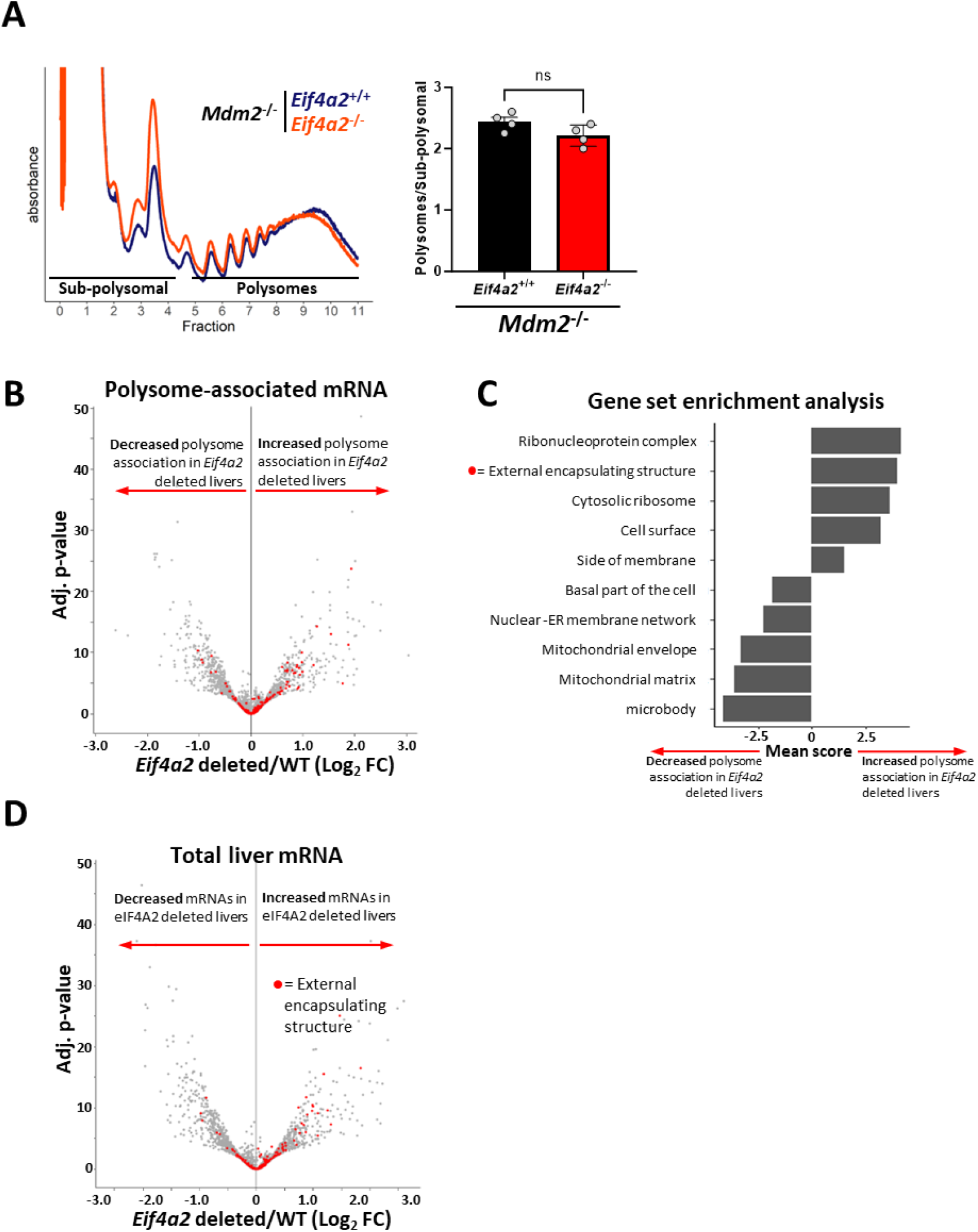
Polysome profiling of wild-type and *Eif4a2* knockout livers. Mice bearing floxed alleles of *Mdm2* (*Mdm2*^fl/fl^) that were either *Eif4a2* WT (*Eif4a2*^WT/WT^) or *Eif4a2* floxed (*Eif4a2*^fl/fl^) were injected with AAV-TBG-Cre (2 X 10^11^ GC per mouse). 4 days following this, livers were homogenized in the presence of cycloheximide and polysomes and sub-polysomes were resolved using sucrose-density gradient centrifugation. The ratio of polysome to sub-polysomes was calculated and is displayed in (**A**). Total (**D**) and polysome-associated (**B, C**) mRNAs were determined by RNAseq. GSEA was performed using the GO term ‘cellular component’ as a reference, and the output from this was ranked according to polysome-associated log2FCs. Redundant terms were collapsed using rrvgo. Terms enriched in genes translationally upregulated following *Eif4a2* deletion have positive scores (**C**). This analysis indicates that mRNAs encoding genes from the external encapsulating structure are enriched in polysomes from *Eif4a2*-deleted livers (**C**) and these are highlighted by red dots in the volcano plots in (**B,D**).

**Fig. S4.**
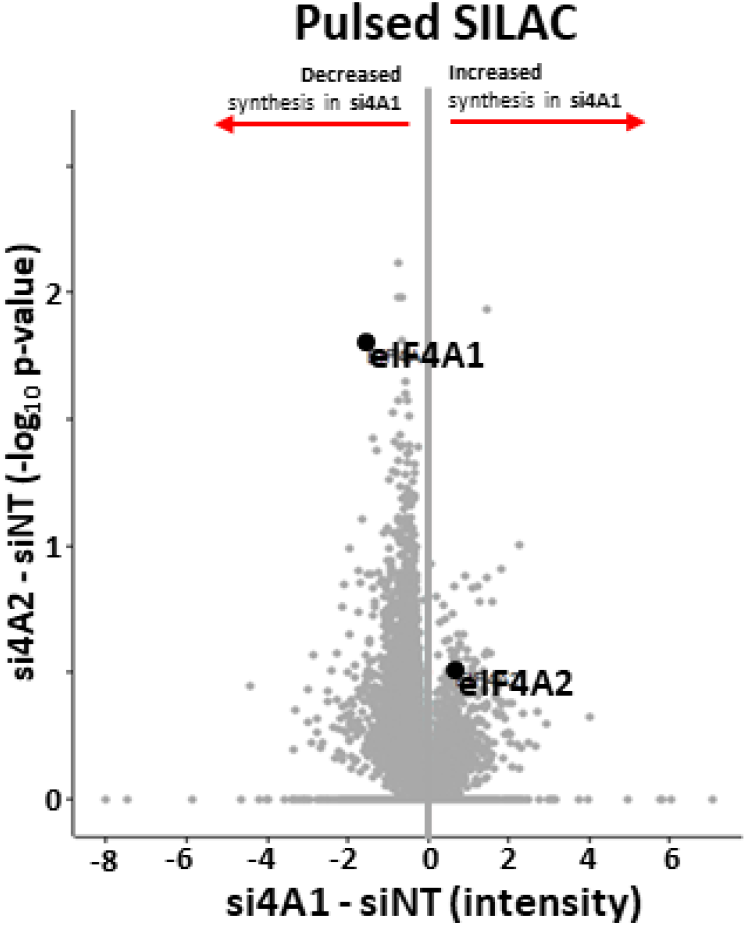
Pulsed-SILAC indicates that eIF4A1 knockdown reduces protein synthesis rates. Fibroblasts were transfected with either non-targeting (siNT) oligonucleotides or those targeting eIF4A1 (si4A1) and were incubated with ‘heavy’ SILAC amino acids (Arg10 & Lys8) for 16h (pulsed-SILAC). Lysates were digested with trypsin and tryptic peptides analysed using mass spectrometry to reveal the influence of eIF4A1 knockdown on the newly synthesized proteome. Data are plotted as differences in intensity (abscissa) and statistical significance (ordinate) between eIF4A1 knockdown and control (si4A1 – siNT).

**Fig. S5.**
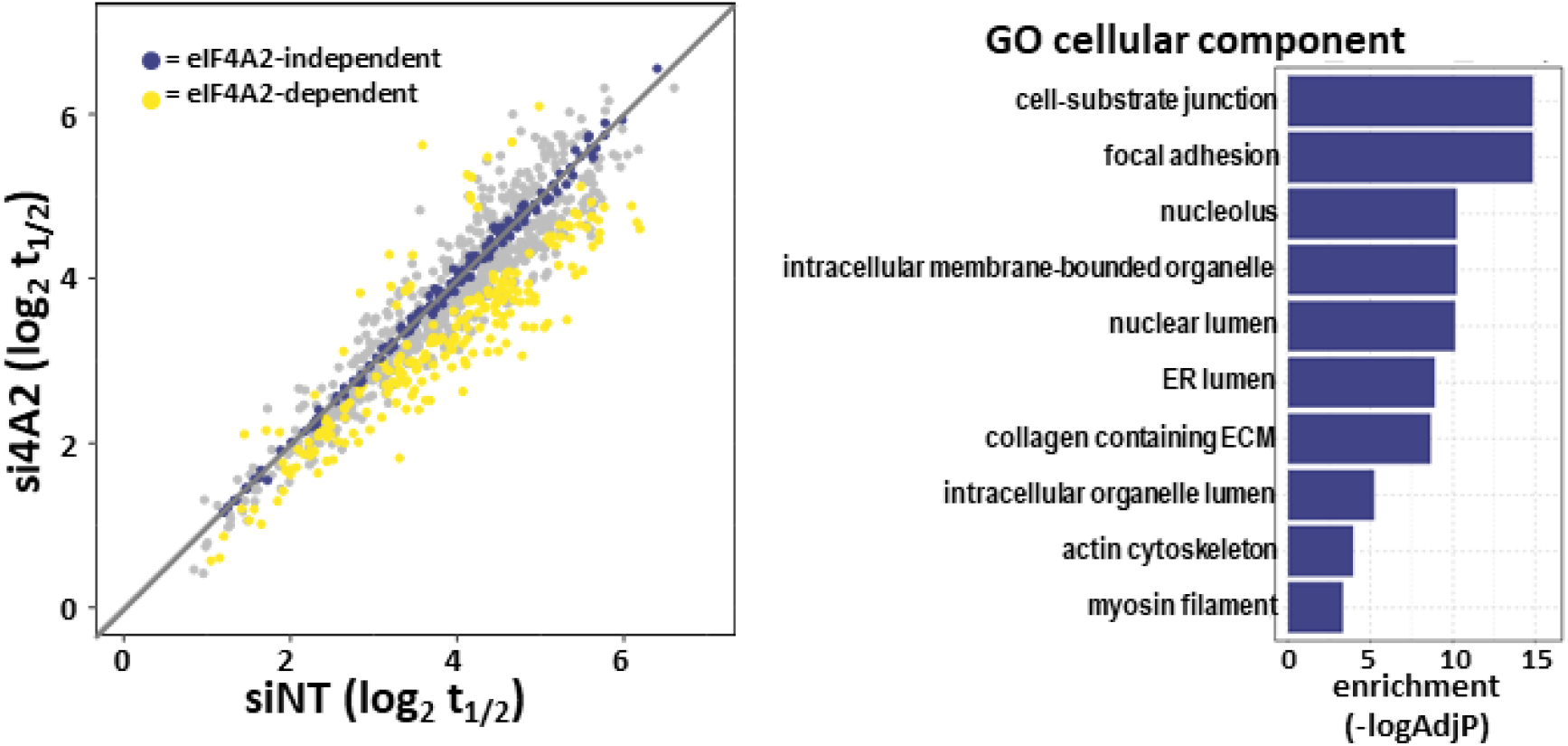
GO terms of proteins whose turnover is increased by eIF4A2 knockdown. Control and eIF4A2 knockdown fibroblasts were incubated with ‘heavy’ SILAC amino acids for the times indicated in Fig. 5A, and tryptic peptides were tandem mass-tagged (TMT) and analysed using mass spectrometry-based proteomics. Turnover rates were derived from the proteomic data and proteins with significantly different t1/2s are denoted using yellow dots (eIF4A2-dependent; left panel). The gene ontology (GO) terms for proteins with significantly different t1/2 are ranked in the right panel.

